# Dual HER2/HER3 Targeting with Antibody-Drug Conjugates Reveals a Novel Therapeutic Strategy for HER2-Mutant Lobular Breast Cancer

**DOI:** 10.64898/2026.06.18.733194

**Authors:** Jie Bin Liu, Ye Cao, Alexander Chih-Chieh Chang, Ryan Jaehne, Daniel D. Brown, Hunter Waltermire, Danielle Tseng, Rinath M. Jeselsohn, Guilherme Nader-Marta, Jagmohan Hooda, Julia Foldi, Marija Balic, Adrian V. Lee, Steffi Oesterreich

## Abstract

Activating HER2 mutations are significantly enriched in both primary and metastatic invasive lobular breast cancer (ILC), with large public datasets of primary breast tumors linking them to a worse prognosis in ILC. Despite their oncogenic role, no FDA-approved therapies currently target HER2-mutant breast cancers. While the HER2-directed antibody-drug conjugate (ADC) trastuzumab deruxtecan (T-DXd) has shown efficacy in HER2-mutant non-small cell lung cancer, its activity in HER2-mutant ILC remains unknown. Using the Caris real-world database, one of the largest cohorts with survival data in advanced breast cancers, we confirmed that HER2 mutations are more prevalent in advanced ILC than in invasive breast cancer of no special type (NST) tumors, are associated with worse survival in both histologies, yet predict improved response to T-DXd across subtypes, highlighting the need for mutation-directed, histology-informed therapies.

Using endogenous HER2-mutant ILC cell lines (UACC3133-S310F, BCK4-L755S) and CRISPR-engineered isogenic ILC models with clinically relevant HER2 mutations (S310F, V777L), we found these mutations drive HER2/HER3 hyperactivation and downstream signaling, conferring increased sensitivity to HER2 tyrosine kinase inhibitors (TKIs) and T-DXd. Mechanistically, HER2 mutants showed enhanced receptor ubiquitination, internalization, and lysosomal degradation upon T-DXd treatment, explaining the observed drug sensitivity. While combining T-DXd with neratinib or the HSP90 inhibitor ganetespib yielded synergistic effects in long-term growth assays, accompanied by increased HER2 ubiquitination, the concurrent hyperactivation of HER3 in HER2-mutant cells suggested that co-targeting HER3 could provide an effective alternative strategy. Accordingly, HER2-mutant ILC exhibited enhanced sensitivity to the HER3-directed ADC patritumab deruxtecan (P-DXd) or LJM716, a HER3-targeting antibody. We further uncovered a previously unrecognized mechanism of P-DXd beyond HER3 ligand blockade and payload delivery: P-DXd promotes HER2/HER3 association, increases HER2 ubiquitination, and enhances T-DXd internalization, resulting in potent synergy with T-DXd. Mechanistically, we identified HER3 extracellular domains I and II as essential for P-DXd binding and for mediating P-DXd-induced HER2/HER3 association, establishing a structural basis for this activity. *In vivo*, both T-DXd and P-DXd suppressed UACC3133 and BCK4 xenograft growth, with combination therapy trending toward greater efficacy and prevented regrowth of tumors.

Extending these findings beyond HER2-mutant ILC, combination treatment with T-DXd and P-DXd demonstrated synergistic activity across multiple breast cancer models, including (i) HER2-amplified NST patient-derived organoids (PDOs) harboring hotspot HER2 mutations, (ii) HER2-wild-type NST PDOs with clinically intrinsic or acquired T-DXd resistance, and (iii) isogenic HER2-mutant ILC PDOs with experimentally induced resistance after prolonged T-DXd exposure. Collectively, these findings support HER2 as an actionable target in HER2-mutant ILC and position T-DXd-based regimens, particularly in combination with HER3 inhibition, as a promising therapeutic strategy for this underserved patient population.

## Introduction

Invasive lobular carcinoma (ILC) is the most common special type of breast cancer with distinctive biology and clinical behavior, representing up to 15% of diagnosed invasive breast cancers [1]. Although most ILC is estrogen receptor positive (ER+) and historically treated according to paradigms for invasive carcinoma of no special type (NST), outcomes remain heterogeneous, with long-term outcome being worse for patients with ILC compared to NST, and precision strategies lag behind biology [2–4]. Genomic studies show that activating HER2 mutations are enriched in ER+/HER2-nonamplified ILC and associated with inferior prognosis compared with wild-type cases [5, 6]. Despite this burden, there are no FDA-approved, mutation-specific therapies for HER2-mutant ILC.

Activating HER2 mutations promote constitutive HER2 signaling and amplify crosstalk with HER3 and EGFR, engaging PI3K/AKT and MAPK/ERK pathways that sustain proliferation and survival [7–9]. Prior mechanistic work has largely relied on lentiviral cDNA overexpression, which can artifactually inflate surface density, distort signaling output, and induce nonphysiologic trafficking [10, 11]. Here, we leverage CRISPR/Cas9-mediated knock-in engineering with matched isogenic controls to introduce clinically observed HER2 mutations under native promoter control. This preserves physiologic regulation and enables direct linkage of mutation-conditioned receptor fate to drug performance. Prior work provides proof-of-concept that knock-in HER2-mutant ILC models are vulnerable to HER2-directed tyrosine kinase inhibitors (TKIs), including poziotinib [12].

Antibody-drug conjugates (ADC) offer a targeted therapeutic approach to target HER2 [13–15]. Trastuzumab deruxtecan (T-DXd), a humanized anti-HER2 antibody linked to a topoisomerase I inhibitor, has shown notable activity in HER2-mutant tumors, but its efficacy in HER2-mutant ILC remains unknown [16]. Crucially, T-DXd performance depends not only on antigen abundance but also on receptor trafficking. Internalization, endosomal routing, and lysosomal processing together determine how efficiently the payload is delivered [15]. Emerging data indicate that some HER2 mutants accelerate receptor internalization and turnover, potentially increasing DXd exposure [10, 11, 17]. Nonetheless, variability in response depth, emergence of resistance, and toxicities highlight the need for strategies that enhance efficacy and extend durability while preserving safety.

These considerations motivate two practical avenues to potentiate activity of T-DXd. First, pharmacologic destabilization of the HER2 complex may increase receptor turnover and ADC processing. The irreversible pan-HER TKI neratinib and the HSP90 inhibitor ganetespib each suppress HER2 signaling and promote receptor degradation in preclinical systems, and TKI/HSP90i + ADC combinations have enhanced internalization and activity in other HER2-driven contexts [18–20]. Their impact alongside T-DXd in HER2-mutant ILC, however, remains to be systematically defined. Second, because HER2 trafficking and signaling are tightly governed by its dimeric state, modulating HER3 offers an orthogonal way to influence both pathway output and T-DXd efficacy. Multiple datasets document elevated HER3 expression in trastuzumab or T-DM1-resistant breast cancers, and ligand-driven HER3 activation can blunt T-DM1 efficacy [21–24]. Clinically tractable HER3-targeted agents, including the HER3-directed ADC patritumab deruxtecan (P-DXd), enable direct testing of this strategy [25–27].

Here, we (i) defined single-agent activity of HER2 TKIs and ADCs across HER2-mutant ILC models, including endogenous and isogenic CRISPR/Cas9 engineered cell lines and patient-derived organoids (PDOs); (ii) tested whether HER2-mutant models exhibit increased receptor ubiquitination and internalization that could mechanistically account for T-DXd efficacy; (iii) evaluated neratinib and ganetespib with T-DXd to determine whether increasing receptor turnover augments T-DXd cytotoxic effects; (iv) and assessed P-DXd alone and combined with T-DXd, extending to dual HER2/HER3 ADC co-targeting *in vitro* and *in vivo*.

## Results

To compare the prevalence of pathogenic or likely pathogenic HER2 mutations in advanced ER+/HER2-nonamplified ILC versus NST, we mapped variant distributions in ILC (n=108) and NST (n=90) (**Figure 1a-b**). Although the overall HER2 mutational landscape broadly overlapped, key hotspots differed between histologies. In ILC, L755S (26.9%), V777L (13.0%), D769Y (10.2%), and S310 substitutions (∼12%) predominated, followed by exon 20 insertions (Y772_A775dup, 6.5%; G778_P780dup, 5.6%). Less frequent events included V842I (5.6%) and L869R (3.7%), with a long tail spanning exons 18-21 and the TM/JM regions. In NST, V777L and L755S each accounted for 21.1% of mutations, followed by G778_P780dup and D769Y (8.9% each), Y772_A775dup (6.7%), and S310F/Y (∼8.8%). Thus, kinase-domain alterations predominated in both histologies, with relative enrichment of L755S and a notable fraction of S310 substitutions in ILC. Consistent with our prior work and that of others [5, 6], HER2 mutations were significantly more frequent in ILC than NST (10.2% versus 2.5%; **Figure 1c**). Domain spectra were broadly similar (**Figure 1d**). Exon analysis showed a dominance of mutations in exons 19 and 20 in both histologies (**Figure 1e**). Survival analysis showed shorter overall survival (OS) in patients with HER2-mutant tumors within each histology (**Figure 1f**). In ILC, median OS was 32.2 months for HER2-mutant versus 41.3 months for HER2-wild-type tumors (HR 1.293; p=0.057); a similar but statistically significant effect was observed in NST (32.0 versus 42.5 months; HR 1.462; p<0.0001). Among HER2-mutant tumors, OS was similar between ILC and NST (median ∼32 months; HR 0.866; p=0.368). In a small treated subcohort that received T-DXd (**Figure 1g**), HER2-mutant cases trended toward improved OS relative to wild-type tumors in both histologies, with median OS of 29.1 versus 16.1 months in ILC (HR 0.580; p=0.17) and 28.4 versus 17.7 months in NST (HR 0.560; p=0.154).

**Figure 1.**
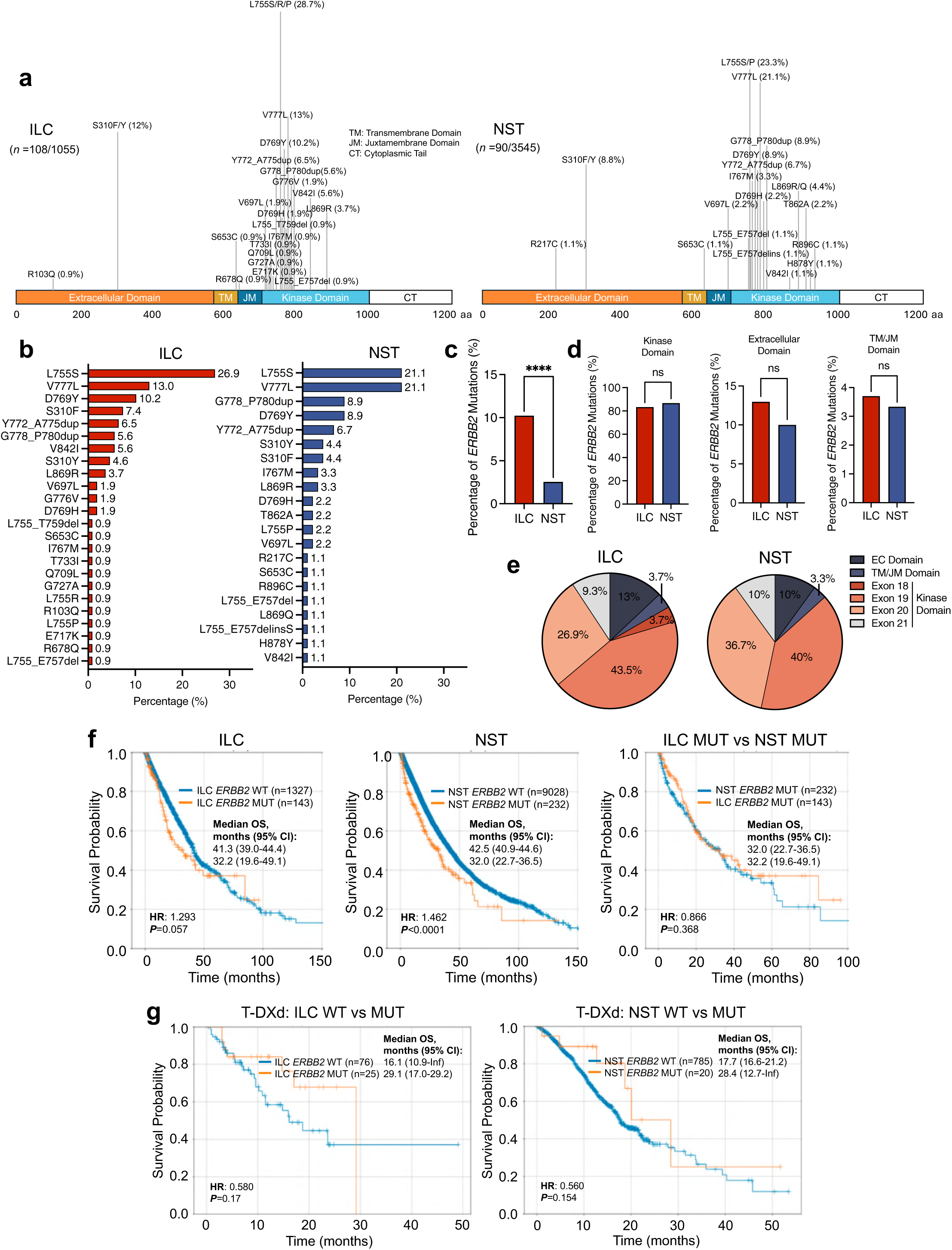
HER2 mutations are enriched in advanced ER⁺/HER2-nonamplified ILC and associate with poor prognosis but improved T-DXd response. (a) Lollipop plot showing the distribution of pathogenic/likely pathogenic HER2 mutations across protein domains. (b) Frequency of pathogenic/likely pathogenic HER2 mutations by histology. (c) Comparison of pathogenic/likely pathogenic HER2 mutation prevalence between ER⁺/HER2-nonamplified ILC and NST. (d) Domain-level comparison of pathogenic/likely pathogenic HER2 mutations between ER⁺/HER2-nonamplified ILC and NST. (e) Exon-level comparison of pathogenic/likely pathogenic HER2 mutations between ER⁺/HER2-nonamplified ILC and NST. (f) Kaplan-Meier overall survival (measured from last clinical contact) comparing ER⁺/HER2-nonamplified ILC and NST breast cancers, stratified by HER2 mutation status. (g) Overall survival among patients with ER⁺/HER2-nonamplified tumors treated with T-DXd in any line of therapy, comparing HER2-mutant and HER2-wild-type cases within each histologic subtype. Statistical significance was determined using the log-rank test. HR, hazard ratio; CI, confidence interval.

To define the functional impact of HER2 mutations in ILC, we generated and curated a panel of CRISPR-engineered knock-in and endogenous HER2-mutant cell line and PDO models (**Extended Data 1a-b**). Using CRISPR/Cas9 prime editing, we introduced the clinically observed activating variants S310F and V777L at the endogenous *ERBB2* locus in two ER+/HER2-nonamplified metastatic ILC cell lines, MDA-MB-134 (hereafter, MM134) and SUM44PE, and in the metastatic ER-/HER2-nonamplified ILC PDO IPM-BO-053. Multiple attempts to generate L755S knock-ins were unsuccessful. In parallel, whole exome sequencing of 17 ILC lines and our breast cancer PDO resource (n=111) identified endogenous HER2-mutant ILC models, including ER+/HER2-nonamplified cell lines UACC3133 (S310F) and BCK4 (L755S) and the primary ER-/HER2-nonamplified PDO IPM-BO-030 (G815R), which harbors a rare activating variant previously described in breast, lung, and hepatobiliary cancers and shown to be sensitive to HER2-selective inhibitors such as zongertinib and sevabertinib [28–32]. Because UACC3133 and BCK4 express higher baseline HER2, we included the HER2-high/wild-type, ER- ILC lines IPH926 and HCC2185 as expression-matched controls, along with the HER2-low/wild-type, ER+ ILC lines SUM44PE and MM134, to reduce confounding effects from receptor abundance. Primary ER+ PDOs IPM-BO-086 and IPM-BO-046 served as HER2-wild-type controls for IPM-BO-030. Sanger sequencing confirmed the S310F and V777L edits and verified endogenous mutations (**Extended Data 1c**). Cellular morphology differed between isogenic wild-type and mutant models (**Extended Data 1d**). In MM134, S310F and V777L HER2 mutants displayed less well-defined cell-cell borders and appeared more cohesive than the more isolated wild-type cells. In SUM44PE, V777L HER2 mutant cultures formed larger, often irregular colonies. In IPM-BO-053 PDO, S310F and V777L HER2 mutants generated larger organoids than wild type. These features are consistent with mutation-driven HER2 pathway hyperactivation and suggest increased proliferative fitness at physiologic expression. We profiled HER2 signaling at baseline and after heregulin (HRG) stimulation (**Extended Data 1e-g, Supplementary Figure 1a-d**). Among endogenous mutants, UACC3133 and BCK4 displayed higher pHER2 and pHER3 at baseline and with ligand, while downstream AKT and ERK phosphorylation was comparable to HER2-high wild-type lines IPH926 and HCC2185 (**Extended Data 1e**). In IPM-BO-030, HRG stimulation produced greater increases in pHER2, pHER3, pAKT, and pERK, consistent with an activating effect of G815R (**Extended Data 1f**). In isogenic models, mutant clones showed higher pHER2, pHER3, pAKT, and pERK following HRG; V777L HER2 mutants also exhibited elevated baseline HER2/HER3 phosphorylation relative to wild-type (**Extended Data 1g**).

To test whether activating HER2 mutations sensitize ILC models to HER2-directed therapy, we performed dose-response assays in isogenic and endogenous models with HER2 TKIs and ADCs. In isogenic MM134 and IPM-BO-053, S310F or V777L HER2 mutants uniformly lowered IC50 values versus wild-type (**Figure 2a, c**). Sensitization was most pronounced for irreversible TKIs and both ADCs, whereas tucatinib showed no consistent shift. These data indicate that physiologic expression of HER2 mutation increases responsiveness across multiple HER2-targeted modalities. Despite increased signaling, HER2-mutant SUM44PE was preferentially sensitive to irreversible TKIs but not to HER2-targeted ADCs (**Supplementary Figure 2a-b**). Transcriptomics indicated elevated efflux/detox genes in SUM44PE that might blunt ADC activity (**Supplementary Figure 2c**). Given this intrinsic resistance signature, SUM44PE was deprioritized for subsequent ADC therapeutic efficacy studies. In endogenous HER2-mutant ILC lines, a similar pattern emerged, with greater sensitivity to T-DXd and TKIs than in HER2 wild-type comparators (**Figure 2b-c**). The HER2-high wild-type line IPH926 also showed some TKI sensitivity. Unlike the isogenic models, the mutant lines were additionally more responsive to tucatinib, indicating model-dependent inhibitor profiles. We tested whether mutant-driven activation yields stronger pharmacodynamic responses by profiling signaling after T-DXd or neratinib under serum starvation and with HRG (**Figure 2d**). In isogenic models, HRG induced greater pHER2/pHER3 and downstream pAKT/pERK in mutants, which were effectively blocked by T-DXd or neratinib. Endogenous mutants showed the same pattern; both agents suppressed pHER2 /pHER3 and downstream nodes under both conditions. As a functional readout of payload delivery, T-DXd increased pH2AX in MM134 and IPM-BO-053, with larger fold changes in S310F and V777L HER2 mutants than in wild-type (**Figure 2e**). UACC3133 and BCK4 showed strong pH2AX induction, whereas HER2-high wild-type IPH926 and HCC2185 increased more modestly.

**Figure 2.**
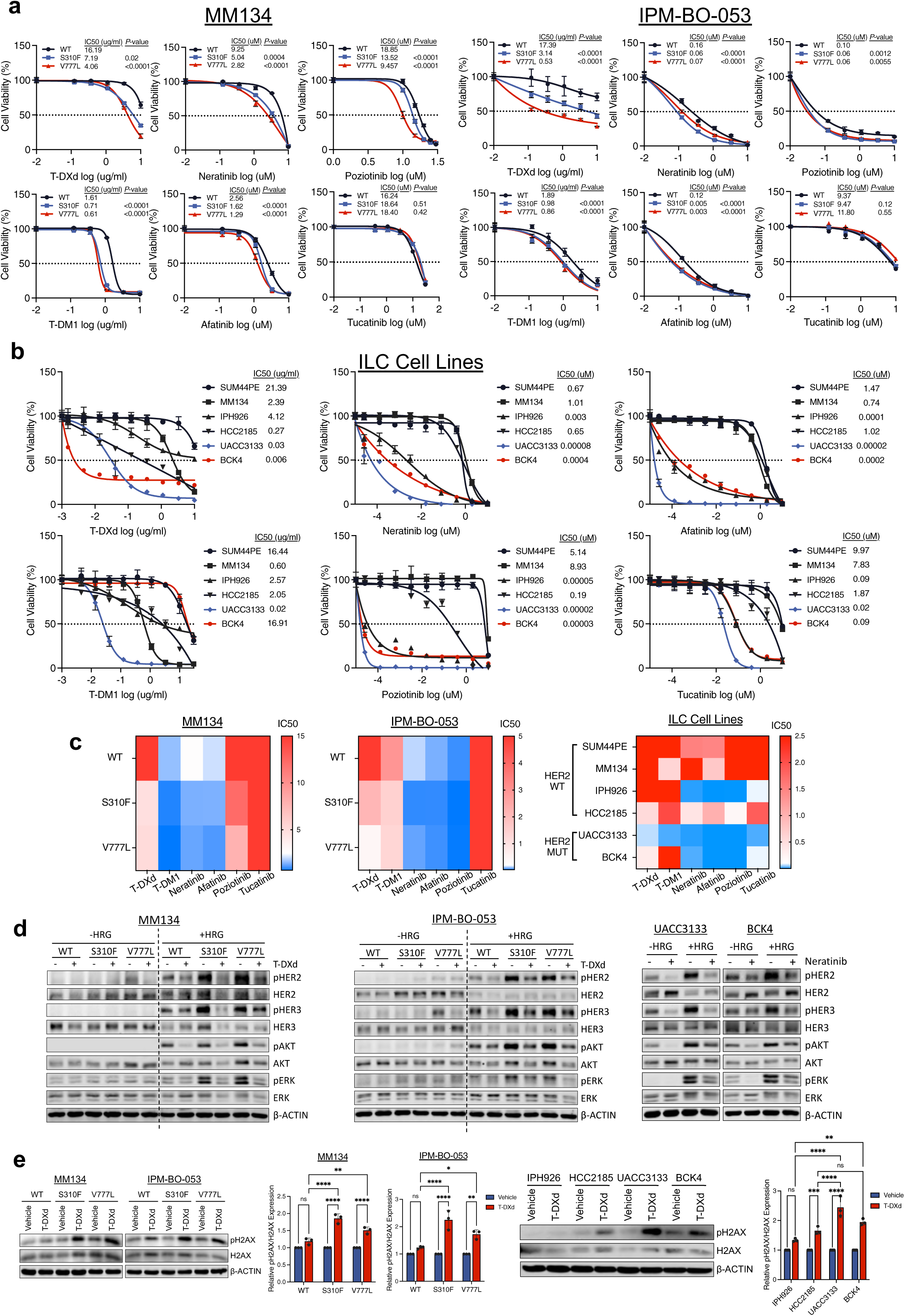
Activating HER2 mutations confer increased sensitivity to HER2-targeted TKIs and ADCs in ILC. (a) Dose-response curves comparing sensitivity of isogenic MM134 and IPM-BO-053 models to HER2-targeted tyrosine kinase inhibitors (TKIs: neratinib, afatinib, poziotinib, tucatinib) and antibody-drug conjugates (ADCs: T-DM1, T-DXd). Error bars represent mean ± SD. (b) Dose-response curves comparing sensitivity of endogenous HER2-mutant (UACC3133, BCK4) and HER2-wild-type (SUM44PE, MM134, IPH926, HCC2185) ILC cell lines to the indicated HER2 TKIs and ADCs. Error bars represent mean ± SD. (c) Heatmap summarizing IC50 values for each agent across isogenic and endogenous models. (d) Immunoblot analysis of HER2/HER3 and downstream signaling in serum-starved isogenic and endogenous models pretreated with T-DXd (10 µg/mL, 4 h) or neratinib (20 nM, 4 h), followed by vehicle or HRG (1 nM) stimulation for 10 or 30 minutes (as indicated). (e) Immunoblot analysis of pH2AX/H2AX ratios following T-DXd treatment (1 µg/mL, 3 days) in isogenic and endogenous models, with quantification normalized to vehicle-treated controls. Error bars represent mean ± SEM. Representative results are shown from three independent experiments (n = 3). Statistical significance was determined using a sum-of-squares Global F-test (IC50 comparisons) or two-way ANOVA; *p* < 0.05.

To determine whether activating HER2 mutations alter receptor turnover and thereby influence T-DXd efficacy, we quantified ubiquitination, internalization, and degradation after HRG or T-DXd across isogenic and endogenous ILC models. HRG markedly increased HER2 ubiquitination in endogenous mutants versus wild-types, with higher mutant basal ubiquitination that further rose with ligand (**Extended Data 2a**). Isogenic mutants recapitulated stronger HRG-induced ubiquitination in S310F and V777L HER2 mutants than in wild-type (**Extended Data 2b**). Cycloheximide assays showed faster HER2 degradation in mutants (**Extended Data 2c**). Bafilomycin A1 and MG132 stabilized HER2, implicating lysosomal and proteasomal pathways, respectively, in HRG-driven turnover (**Extended Data 2d**). Building on this, we determined whether mutations also enhance T-DXD/HER2 complex internalization. Live-cell imaging using pHrodo-T-DXd showed markedly higher T-DXD/HER2 uptake in HER2-mutant lines than in wild-type comparators (**Figure 3a**). Isogenic models mirrored these trends, where SUM44PE and IPM-BO-053 mutants accumulated more pHrodo-T-DXd than wild-type (**Figure 3b-c**). SUM44PE was prioritized over MM134 for internalization assays owing to higher baseline HER2, enabling robust fluorescence quantification. To test whether enhanced T-DXd/HER2 uptake in HER2-mutant models requires active endocytosis, we inhibited dynamin-mediated vesicle scission with dyngo-4a. Dyngo-4a pretreatment markedly reduced T-DXd internalization across all models (**Supplementary Figure 3a**). Concordantly, dyngo-4a impaired T-DXd cytotoxicity, increasing IC50 values in UACC3133 and BCK4 (**Supplementary Figure 3b**). These results support a dynamin-dependent mechanism linking receptor trafficking to T-DXd response.

**Figure 3.**
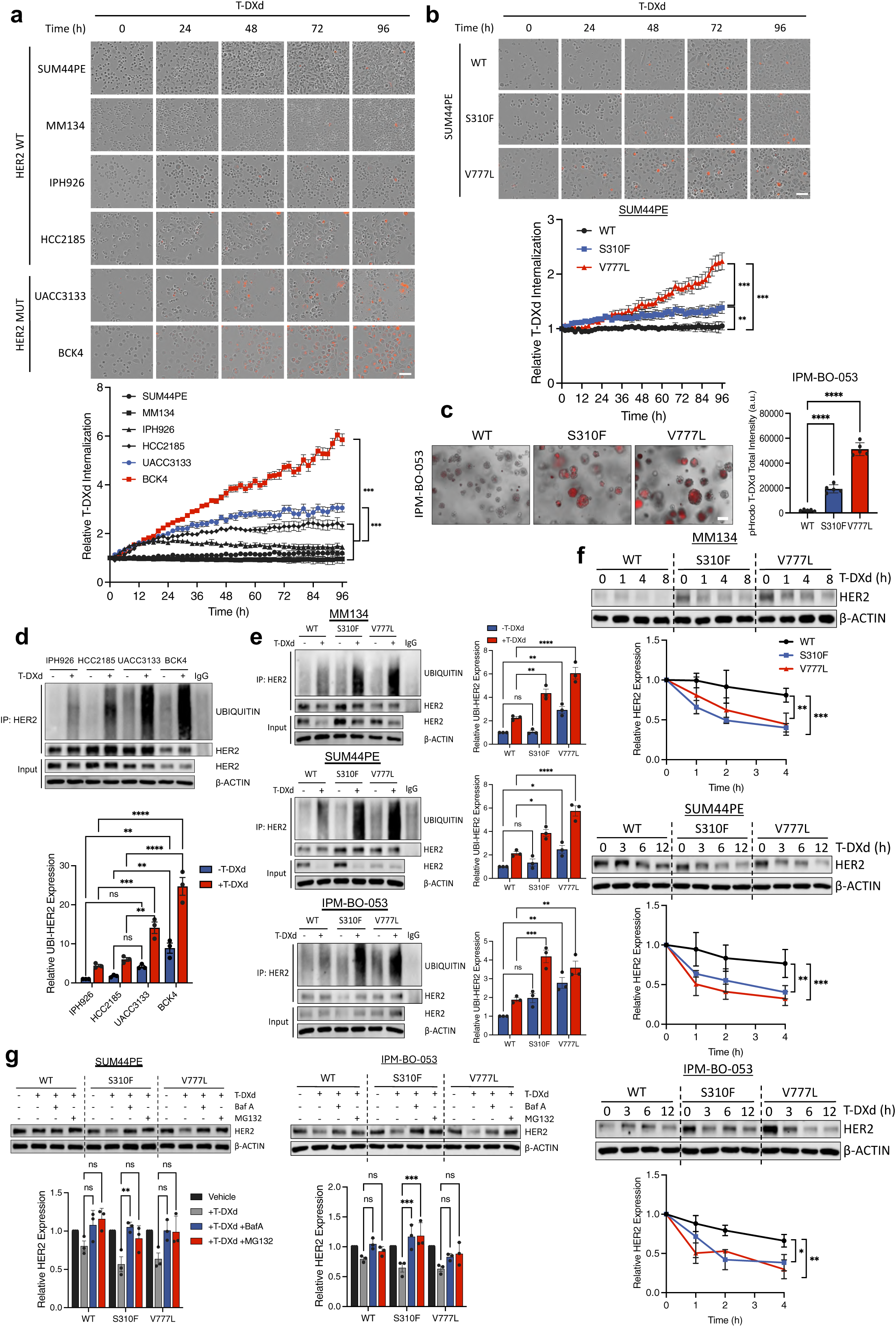
HER2-mutant ILC cells show increased HER2 ubiquitination, internalization, and lysosomal degradation upon T-DXd. (a) Live-cell imaging of pHrodo-T-DXd (1 µg/mL) internalization in endogenous HER2-mutant (UACC3133, BCK4) and HER2-wild-type ILC cell lines (SUM44PE, MM134, IPH926, HCC2185), with quantification of pHrodo fluorescence intensity over time normalized to baseline (t = 0) and confluency. Error bars represent mean ± SD. (b) Live-cell imaging of pHrodo-T-DXd (1 µg/mL) internalization in isogenic SUM44PE models, with quantification normalized to baseline (t = 0) and confluency. Error bars represent mean ± SD. (c) Live-cell imaging of pHrodo-T-DXd (1 µg/mL) internalization in isogenic IPM-BO-053 models at 72 hours, with quantification of pHrodo fluorescence intensity normalized to LysoBrite™ Green (10 nM)-labeled lysosomes. Error bars represent mean ± SD. (d) Immunoblot analysis of HER2 ubiquitination in serum-starved endogenous HER2-mutant (UACC3133, BCK4) and HER2-wild-type (IPH926, HCC2185) ILC cell lines pretreated with bafilomycin A1 (Baf A1; 100 nM, 2 h) followed by vehicle or T-DXd (1 µg/mL, 4 h), with quantification normalized to vehicle-treated IPH926. Error bars represent mean ± SEM. (e) Immunoblot analysis of HER2 ubiquitination in serum-starved isogenic MM134, SUM44PE, and IPM-BO-053 models pretreated with Baf A1 (100 nM, 2 h) followed by vehicle or T-DXd (1 µg/mL, 4 h), with quantification normalized to vehicle-treated wild-type controls. Error bars represent mean ± SEM. (f) Cycloheximide chase assay assessing HER2 degradation kinetics in isogenic models following T-DXd (1 µg/mL) stimulation, with HER2 expression normalized to baseline (t = 0). Error bars represent mean ± SEM. (g) Pharmacologic stabilization assays in isogenic models pretreated with lysosomal inhibitor Baf A1 (100 nM, 2 h) or proteasomal inhibitor MG132 (20 µM, 2 h), followed by T-DXd (1 µg/mL, 4 h), with HER2 expression normalized to vehicle. Error bars represent mean ± SEM. Representative results are shown from three independent experiments (n = 3). Statistical significance was determined using two-way ANOVA; *p* < 0.05.

Having established accelerated T-DXd/HER2 internalization in HER2-mutant ILC cells, we tested whether increased drug-induced receptor ubiquitination underlies this behavior. T-DXd markedly increased HER2 ubiquitination, with the largest effects in UACC3133 and BCK4 (**Figure 3d**). Isogenic MM134, SUM44PE, and IPM-BO-053 recapitulated this with stronger ubiquitination in mutants than in wild-type (**Figure 3e**). Cycloheximide assays showed faster HER2 degradation in mutants following T-DXd (**Figure 3f**), whereas bafilomycin A1 and MG132 stabilized HER2 in SUM44PE and IPM-BO-053, implicating lysosomal and proteasomal pathways, consistent with HRG-driven turnover (**Figure 3g**). Lastly, we visualized lysosomal routing of T-DXd using live-cell colocalization of pHrodo-T-DXd with LysoBrite Green (**Extended Data 3**). Both isogenic and endogenous HER2-mutant models showed stronger pHrodo-T-DXd signal within LysoBrite-positive areas than wild-types. These data indicate that activating HER2 mutations accelerate HER2 ubiquitination, internalization, and lysosomal degradation, thereby enhancing T-DXd delivery.

As T-DXd efficacy is dependent upon HER2 turnover, we assessed whether low-dose neratinib or ganetespib would augment T-DXd-induced HER2 ubiquitination. In isogenic and endogenous models, T-DXd alone increased ubiquitination, the low-dose treatment with neratinib or ganetespib alone had little effect, and co-treatment further amplified ubiquitination beyond either agent (**Figure 4a-b**). Thus, mild HER2 kinase or HSP90 inhibition boosts T-DXd-driven HER2 ubiquitination. We next tested whether pharmacologic destabilization deepens pathway blockade and augments T-DXd-induced DNA damage. In isogenic models, HRG increased pHER2 and pHER3 with concordant AKT and ERK activation; T-DXd or neratinib each suppressed this activation, and the combination produced the most complete inhibition, especially in HER2 mutants (**Figure 4c-d**). Endogenous models showed the same pattern, with combinations outperforming single agents (**Figure 4e-f**). Ganetespib behaved similarly; T-DXd plus ganetespib suppressed signaling more than either agent alone across models. pH2AX induction was minimal with neratinib or ganetespib alone, markedly increased with T-DXd, and further increased with either combination (**Figure 4g-i**). The extent of pH2AX tracked with upstream suppression, consistent with accelerated receptor turnover and more efficient T-DXd payload delivery. We next evaluated whether signaling blockade and DNA damage translated into growth control. In short-term assays across endogenous and isogenic models, combining low-dose neratinib or ganetespib with T-DXd produced mixed effects, ranging from additive to synergistic growth suppression in select contexts (**Figure 4j**). Because DXd is most effective in S phase and ILC cells proliferate slowly, we extended studies in UACC3133 and BCK4 to 3-4-week clonogenic assays. In contrast to the short-term results, both agents synergized with T-DXd, producing pronounced colony inhibition (**Figure 4k**).

**Figure 4.**
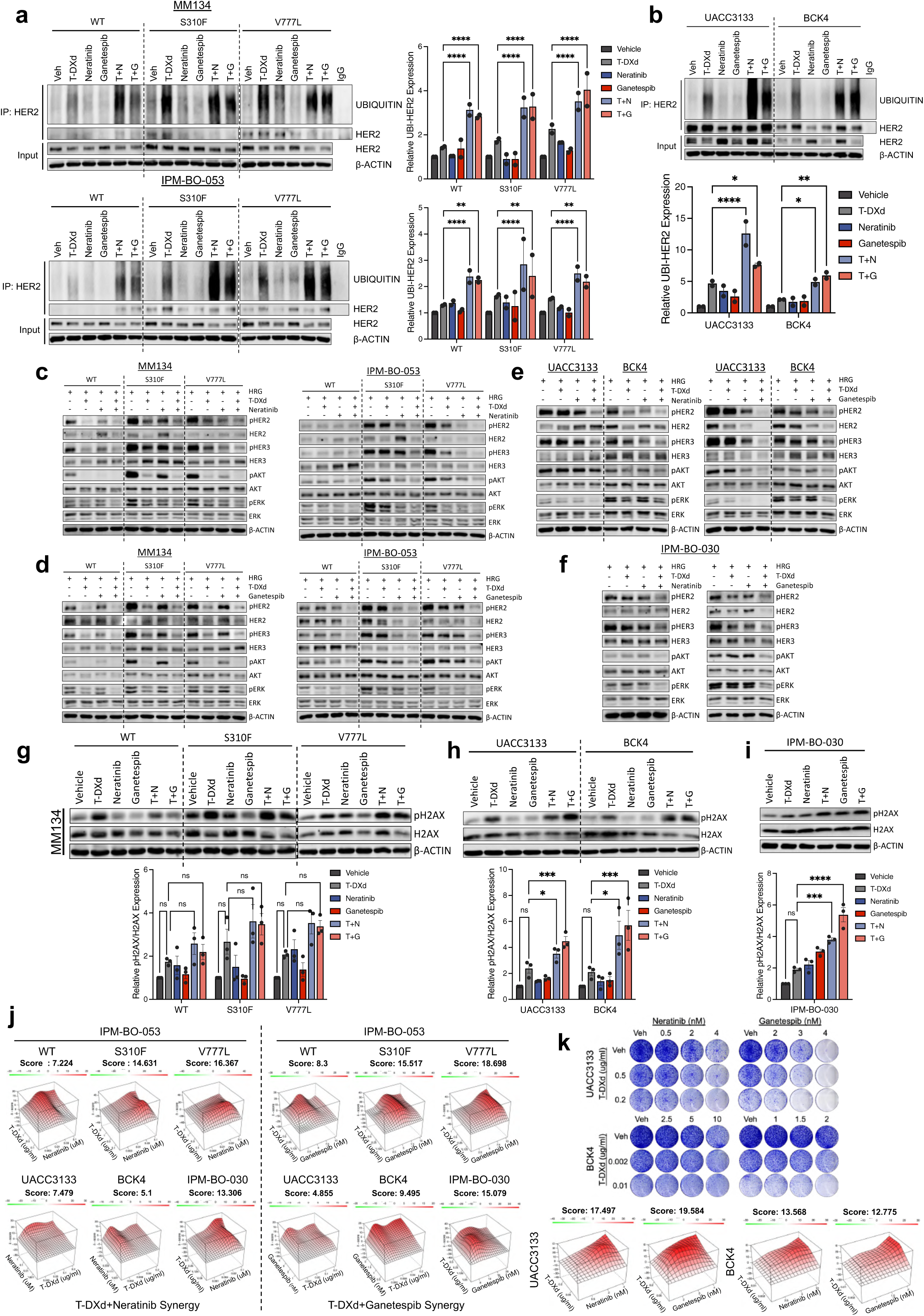
Neratinib or ganetespib induces HER2 ubiquitination and enhances T-DXd efficacy in HER2-mutant ILC. (a) Immunoblot analysis of HER2 ubiquitination in serum-starved isogenic MM134 and IPM-BO-053 models pretreated with bafilomycin A1 (Baf A1; 100 nM, 2 h) followed by T-DXd (1 µg/mL), neratinib (20 nM), ganetespib (20 nM), or the indicated combinations (4 h), with quantification normalized to wild-type controls. Error bars represent mean ± SEM. (b) Immunoblot analysis of HER2 ubiquitination in serum-starved UACC3133 and BCK4 pretreated with Baf A1 (100 nM, 2 h) followed by T-DXd (1 µg/mL), neratinib (20 nM), ganetespib (20 nM), or the indicated combinations (4 h), with quantification normalized to vehicle-treated controls. Error bars represent mean ± SEM. (c) Immunoblot analysis of HER2/HER3 and downstream signaling in serum-starved isogenic MM134 and IPM-BO-053 models pretreated with T-DXd (1 µg/mL, 4 h), neratinib (10 nM, 4 h), or the combination, followed by vehicle or HRG (1 nM) stimulation for 10 or 30 minutes, respectively. (d) Immunoblot analysis of HER2/HER3 and downstream signaling in serum-starved isogenic MM134 and IPM-BO-053 models pretreated with T-DXd (1 µg/mL, 4 h), ganetespib (7.5 nM, 4 h), or the combination, followed by vehicle or HRG (1 nM) stimulation for 10 or 30 minutes, respectively. (e) Immunoblot analysis of HER2/HER3 and downstream signaling in serum-starved UACC3133 and BCK4 pretreated with T-DXd (1 µg/mL, 4 h), neratinib (10 nM, 4 h), ganetespib (7.5 nM, 4 h), or the indicated combinations, followed by vehicle or HRG (1 nM) stimulation for 10 minutes. (f) Immunoblot analysis of HER2/HER3 and downstream signaling in serum-starved IPM-BO-030 pretreated with T-DXd (1 µg/mL, 4 h), neratinib (10 nM, 4 h), ganetespib (7.5 nM, 4 h), or the indicated combinations, followed by vehicle or HRG (1 nM) stimulation for 10 minutes. (g-i) Immunoblot analysis of pH2AX/H2AX ratios following 72-hour treatment with T-DXd (1 µg/mL), neratinib (10 nM), ganetespib (7.5 nM), or the indicated combinations in isogenic MM134 models (g), UACC3133 and BCK4 (h), and IPM-BO-030 (i). Error bars represent mean ± SEM. (j) Short-term (7-day) proliferation analysis of T-DXd, neratinib, ganetespib, and combinations in endogenous models (UACC3133, BCK4, IPM-BO-030) and isogenic IPM-BO-053 models. (k) Long-term (3-4 week) clonogenic survival analysis of T-DXd, neratinib, ganetespib, and combinations in UACC3133 and BCK4). Synergy scores were calculated using the HSA model; values > 10 indicate synergy. Representative results are shown from three independent experiments (n = 3). Statistical significance was determined using one-way or two-way ANOVA; *p* < 0.05.

Signaling analyses across our endogenous and isogenic HER2-mutant ILC models showed disproportionate HER3 activation relative to wild-type, nominating HER3 as a potential co-driver. To evaluate clinical relevance, we compared ER+/HER2-mutant ILC with NST across TCGA, METABRIC, and SCAN-B (**Figure 5a**). In all three RNA-seq cohorts, a signature score of HER3 activity was higher in ILC [33], reaching statistical significance in SCAN-B. Protein data from TCGA RPPA were concordant, with pHER3 trending higher in HER2-mutant ILC than in NST while total HER3 remained similar between subtypes (**Figure 5b**). The mismatch between phosphorylated and total HER3 indicates preferential activation rather than overexpression in HER2-mutant ILC, suggesting subtype-specific rewiring of receptor signaling. We tested dependence on HER2/HER3 co-signaling in HER2-mutant ILC using siRNA to deplete HER2, HER3, or both in endogenous HER2-wild-type and mutant lines (**Supplementary Figure 4a-b**). HER2 knockdown markedly impaired proliferation in UACC3133 and BCK4, with only modest effects in wild-type controls. HER3 silencing reduced viability to a similar extent, and dual knockdown produced the strongest growth inhibition, most pronounced in mutants. These findings indicate co-dependence on HER2 and HER3 for sustained proliferation in HER2-mutant ILC.

**Figure 5.**
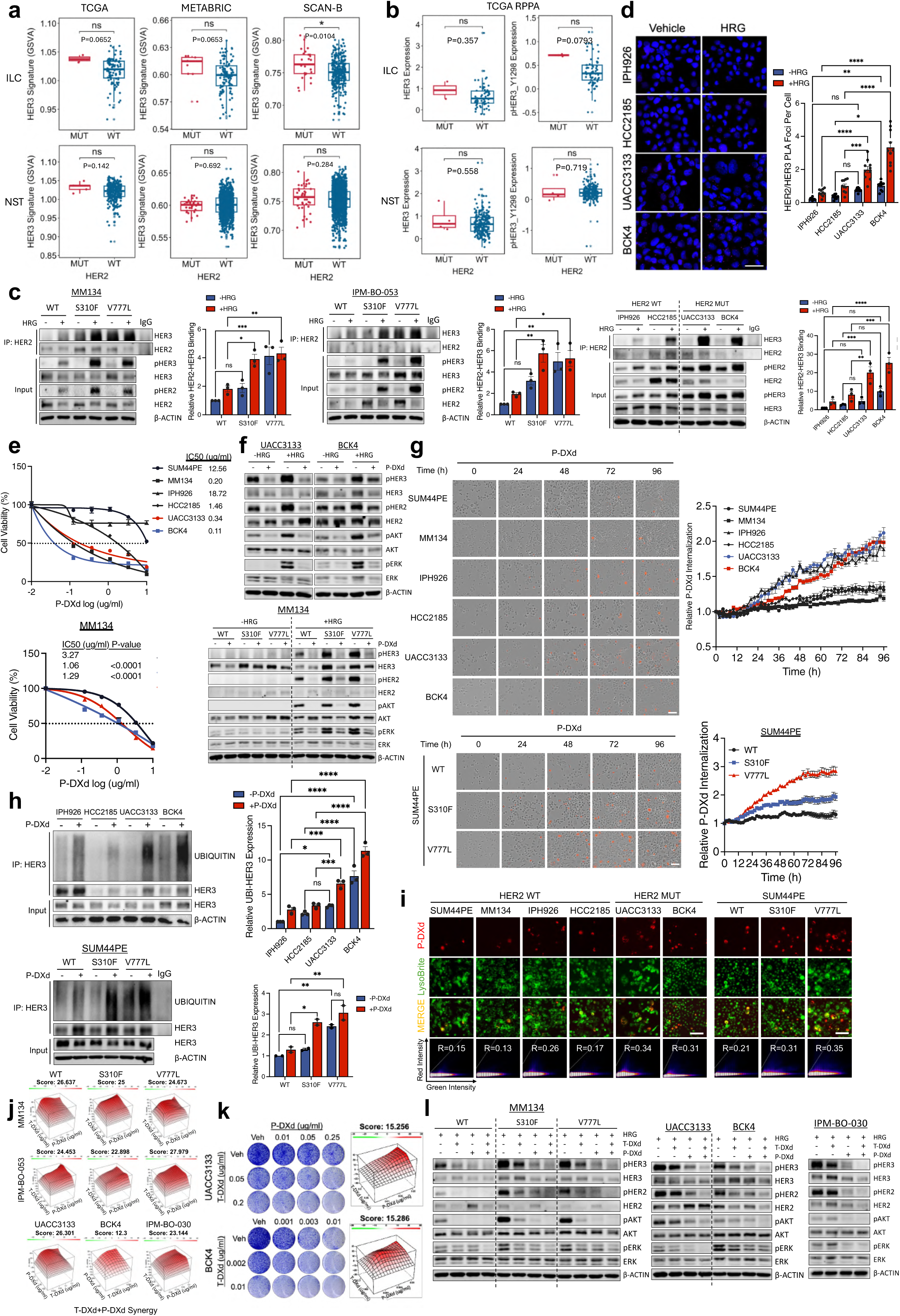
Inhibition of HER3 by P-DXd synergizes with T-DXd. (a) HER3 signaling signature score analysis of bulk RNA-seq data from TCGA, METABRIC, and SCAN-B cohorts comparing ER⁺/HER2-mutant ILC and NST breast cancers. (b) Reverse-phase protein array (RPPA) analysis of total HER3 and phosphorylated HER3 (pHER3; Y1289) protein levels in the TCGA breast cancer cohort comparing ER⁺/HER2-mutant ILC and NST. (c) Immunoblot analysis of HER2/HER3 association in serum-starved isogenic and endogenous models treated with vehicle or HRG (1 nM, 10 min), with quantification normalized to vehicle-treated wild-type controls. Error bars represent mean ± SEM. (d) Proximity ligation assay (PLA) images of HER2/HER3 interactions in serum-starved endogenous models treated with vehicle or HRG (1 nM, 10 min), with quantification of HER2/HER3 puncta normalized to vehicle-treated IPH926. Error bars represent mean ± SD. (e) Dose-response curves evaluating sensitivity of endogenous and isogenic models to P-DXd. Error bars represent mean ± SD. (f) Immunoblot analysis of HER2/HER3 and downstream signaling in serum-starved endogenous and isogenic models pretreated with P-DXd (10 µg/mL, 4 h) followed by vehicle or HRG (1 nM, 10 min). (g) Live-cell imaging of pHrodo-P-DXd internalization in endogenous and isogenic models, with quantification over time normalized to baseline (t = 0) and confluency. Error bars represent mean ± SD. (h) Immunoblot analysis of HER3 ubiquitination in serum-starved endogenous and isogenic models pretreated with bafilomycin A1 (Baf A1; 100 nM, 2 h) followed by vehicle or P-DXd (1 µg/mL, 4 h), with quantification normalized to vehicle-treated wild-type controls. Error bars represent mean ± SEM. (i) Live-cell fluorescence imaging of pHrodo-P-DXd (1 µg/mL; red) and LysoBrite™ Green-labeled lysosomes (10 nM) after 48 hours; co-localization was quantified using Pearson’s correlation coefficient. (j) Short-term (7-day) proliferation analysis of T-DXd, P-DXd, and combinations in isogenic and endogenous models. (k) Long-term (3-4 week) clonogenic survival analysis of T-DXd, P-DXd, and combinations in UACC3133 and BCK4. Synergy scores were calculated using the HSA model; values > 10 indicate synergy. (l) Immunoblot analysis of HER2/HER3 and downstream signaling in serum-starved isogenic and endogenous models pretreated with T-DXd (1 µg/mL, 4 h), P-DXd (1 µg/mL, 4 h), or the combination, followed by vehicle or HRG (1 nM) stimulation for 10 or 30 minutes. Representative results are shown from three independent experiments (n = 3). Statistical significance was determined using the Mann-Whitney U test or two-way ANOVA; *p* < 0.05.

To probe HER2/HER3 co-activation in HER2-mutant ILC, we assessed HRG-induced HER2/HER3 association (**Figure 5c**). In isogenic models, V777L HER2 mutants showed elevated basal HER2/HER3 association; HRG further increased complex formation more in mutants than in wild-type and was accompanied by higher pHER2 and pHER3. Endogenous mutants UACC3133 and BCK4 showed the same pattern, with greater basal association and additional HRG-induced increases with concordant phosphorylation. Proximity ligation assays confirmed higher baseline HER2/HER3 proximity and a larger HRG-induced rise in foci in mutants (**Figure 5d**). These data indicate that activating HER2 mutations prime the receptor for enhanced HER3 engagement, amplifying HRG-dependent signaling.

Given HER3 co-activation in HER2-mutant ILC, we tested effects of the HER3-directed ADC P-DXd. Endogenous and isogenic HER2 mutants were markedly more sensitive than wild-types (**Figure 5e**). MM134, which was relatively refractory to T-DXd, showed clear P-DXd sensitivity, consistent with higher HER3 expression. We assessed signaling inhibition by P-DXd (**Figure 5f**). Under basal and HRG stimulation, P-DXd reduced pHER3/pHER2 and dampened AKT/ERK phosphorylation in both isogenic models, with consistently stronger inhibition in mutants. Given P-DXd sensitivity, we tested whether HER2 mutations also accelerate P-DXd internalization (**Figure 5g**). Live-cell pHrodo-P-DXd imaging showed faster uptake in UACC3133 and BCK4 than in most wild-type comparators; IPH926 matched mutant-like rates, suggesting additional cell-specific factors. Isogenic SUM44PE mutants likewise internalized P-DXd more rapidly than wild-type. These results extend the T-DXd findings, indicating that HER2 mutations enhance HER3-driven signaling and accelerate P-DXd uptake. To assess endocytic dependence, we inhibited dynamin with dyngo-4a (**Supplementary Figure 5a-b**). Live-cell imaging showed a marked reduction in P-DXd internalization, and cytotoxicity decreased accordingly in UACC3133 and BCK4. Thus, P-DXd activity in HER2-mutant ILC relies on dynamin-mediated endocytosis, paralleling the trafficking dependence observed for T-DXd.

Building on the internalization data, we tested whether faster P-DXd uptake in HER2-mutant models is coupled to enhanced HER3 ubiquitination and lysosomal delivery (**Figure 5h**). In endogenous models, basal HER3 ubiquitination was higher in BCK4 and increased further after P-DXd treatment in both UACC3133 and BCK4, relative to wild-type controls. Isogenic models recapitulated this pattern, with V777L HER2 mutants showing higher baseline ubiquitination that increased with P-DXd, and S310F HER2 mutants exhibiting greater induction than wild-type. LysoBrite imaging confirmed P-DXd colocalization with lysosomes across models, with higher lysosomal accumulation in mutants (**Figure 5i**). Taken together with the uptake data, these findings show that activating HER2 mutations increase HER3 ubiquitination and accelerate P-DXd internalization and lysosomal routing.

Because HER2 mutations accelerate internalization of T-DXd and P-DXd, we tested dual targeting (**Figure 5j-k**). In short-term viability assays, T-DXd plus P-DXd showed consistent synergy across endogenous and isogenic modes; synergy was also evident in HER2-wild-type ILC. In long-term clonogenic assays, the combination synergistically suppressed survival beyond either monotherapy. To probe the basis of T-DXd/P-DXd synergy, we profiled pathway inhibition with each agent alone and in combination across isogenic and endogenous models (**Figure 5k**). HRG increased pHER2/pHER3 and activated AKT/ERK, more strongly in mutants. Either ADC alone partially reduced these signals, whereas dual-ADC treatment produced the strongest suppression. Building on the dual-ADC synergy, we tested a broader panel of HER3-directed agents for single-agent and combinatorial effects with T-DXd (**Extended Data 4a**). LJM716, which locks HER3 in an inactive conformation and blocks both ligand-independent and ligand-dependent activation, showed selective cytotoxicity in HER2-mutant ILC, while most HER2-wild-type lines were resistant, with IPH926 as an exception. Patritumab and pertuzumab showed little activity across models. Zenocutuzumab was potent in UACC3133 but not in BCK4, suggesting an S310F-specific dependency. In endogenous models, LJM716 reduced pHER3 and pHER2 and attenuated AKT and ERK signaling (**Extended Data 4b**). Adding T-DXd further strengthened inhibition of HER2/HER3/AKT phosphorylation, closely matching the dual-ADC profile (**Extended Data 4c-d**). Functionally, LJM716 plus T-DXd was synergistic in short-term viability and long-term clonogenic assays, supporting combined HER2 and HER3 blockade as an effective strategy in HER2-mutant ILC (**Extended Data 4e-f**).

We compared DNA damage accumulation across all T-DXd-based combinations in endogenous and isogenic models (**Extended Data 5a-c**). In UACC3133, BCK4 and IPM-BO-030, T-DXd increased pH2AX, and adding neratinib or ganetespib further amplified the signal, consistent with accelerated HER2 turnover. Dual-ADC treatment showed the same pattern, with P-DXd alone inducing pH2AX and the combination with T-DXd producing a larger increase. Pairing LJM716 with T-DXd likewise elevated pH2AX beyond monotherapy. The isogenic organoids recapitulated these effects in both wild-type and mutant backgrounds. Lastly, we assessed apoptosis across single-agent and combination treatments in isogenic IPM-BO-053 organoids (**Extended Data 5d**). Live-cell caspase-3/7 imaging showed apoptosis with T-DXd that increased further with neratinib, ganetespib, P-DXd, or LJM716, with the largest effects in HER2-mutants. Immunoblotting corroborated this pattern, showing modest PARP cleavage on T-DXd alone and markedly higher cleavage with combinations, again more pronounced in mutants (**Extended Data 5e**). Thus, HER2 destabilization or HER3 co-inhibition amplifies T-DXd-induced apoptosis, underpinning the observed synergy.

To investigate mechanisms of T-DXd and P-DXd synergy beyond accelerated internalization, we examined receptor-level crosstalk during dual engagement. We hypothesized that P-DXd would disrupt HRG-induced HER2/HER3 interactions and block activation. Indeed, P-DXd suppressed phosphorylation of HER2, HER3, AKT, and ERK, confirming blockade of HRG-driven signaling. Unexpectedly, P-DXd increased, rather than disrupted, HER2/HER3 association in both isogenic and endogenous mutants. T-DXd alone minimally affected complex assembly, and adding T-DXd did not diminish the P-DXd-induced association (**Figure 6a-b**). PLA corroborated the co-IP findings, with HRG increasing HER2/HER3 proximity, T-DXd alone showing minimal impact, and P-DXd markedly elevating PLA foci, most prominently in HER2-mutant lines. The T-DXd plus P-DXd combination maintained this enhanced PLA signal (**Figure 6c**).

**Figure 6.**
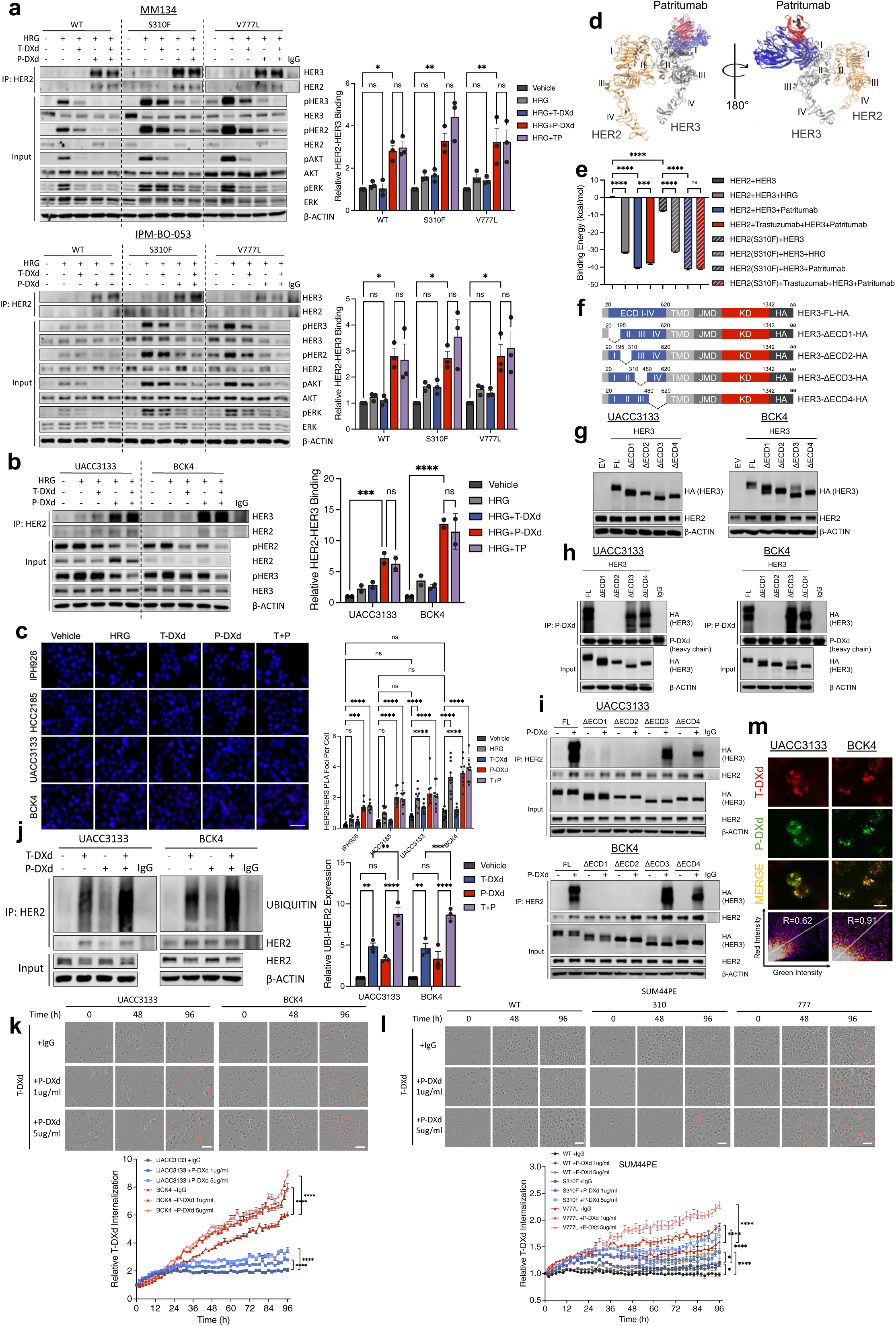
P-DXd induces HER2/HER3 interaction, HER2 ubiquitination, and internalization of T-DXd. (a) Immunoblot analysis of HER2/HER3 association and signaling inhibition in isogenic models treated with vehicle, HRG (1 nM), T-DXd (1 µg/mL), P-DXd (1 µg/mL), or the combination, with quantification normalized to vehicle-treated controls. Error bars represent mean ± SEM. (b) Immunoblot analysis of HER2/HER3 association and signaling inhibition in UACC3133 and BCK4 treated with HRG (1 nM), T-DXd (1 µg/mL), P-DXd (1 µg/mL), or the combination, with quantification normalized to vehicle-treated controls. Error bars represent mean ± SEM. (c) PLA images showing HER2/HER3 interactions in serum-starved HER2-mutant (UACC3133, BCK4) and HER2-wild-type (IPH926, HCC2185) ILC cells treated with vehicle, HRG (10 nM), T-DXd (1 µg/mL), P-DXd (1 µg/mL), or the combination, with quantification of PLA foci per cell normalized to vehicle-treated controls. Error bars represent mean ± SEM. (d) Molecular docking model illustrating the interaction of patritumab with the HER3 extracellular domain. (e) Comparative binding energy analysis of HER2/HER3 complexes bound to HRG or patritumab, with or without trastuzumab co-docking, in wild-type and S310F-mutant HER2 models. Error bars represent mean ± SEM. (f) Schematic representation of full-length (FL) HER3 and truncated constructs lacking individual extracellular domains (ΔECD1-ΔECD4). (g) Immunoblot analysis of UACC3133 and BCK4 cells stably expressing the indicated HER3 constructs. (h) Immunoblot analysis of P-DXd binding to FL and truncated HER3 constructs (ΔECD1-ΔECD4) in UACC3133 and BCK4 using P-DXd as the capturing antibody. (i) Immunoblot analysis of P-DXd-induced HER2/HER3 interactions in UACC3133 and BCK4 expressing FL or truncated HER3 constructs. (j) Immunoblot analysis of HER2 ubiquitination in UACC3133 and BCK4 treated with vehicle, T-DXd (1 µg/mL), P-DXd (1 µg/mL), or the combination, with quantification normalized to vehicle-treated controls. Error bars represent mean ± SEM. (k) Live-cell imaging of pHrodo-T-DXd internalization in UACC3133 and BCK4 treated with T-DXd (1 µg/mL) alone or in combination with P-DXd (1 or 5 µg/mL), with quantification over time normalized to baseline (t = 0) and confluency. Error bars represent mean ± SD. (l) Live-cell imaging of pHrodo-T-DXd internalization in isogenic SUM44PE models treated with T-DXd (1 µg/mL) alone or in combination with P-DXd (1 or 5 µg/mL), with quantification over time normalized to baseline (t = 0) and confluency. Error bars represent mean ± SD. (m) Co-tracking analysis of T-DXd (red) and P-DXd (green) internalization after 48 hours in UACC3133 and BCK4; co-localization was quantified using Pearson’s correlation coefficient. Representative results are shown from three independent experiments (n = 3). Statistical significance was determined using two-way ANOVA; *p* < 0.05.

As the mechanism of binding of patritumab to HER3 is currently unknown, we investigated how P-DXd promotes HER2/HER3 complex formation by molecular modeling of patritumab-HER3 binding. Molecular docking analysis placed patritumab primarily on HER3 ECD1, overlapping the HRG interface (**Figure 6d**). Binding-energy analysis showed HRG strongly stabilizes HER2/HER3, and patritumab confers comparable or greater stabilization in both wild-type and S310F settings (**Figure 6e**). Co-docking with trastuzumab modestly reduced affinity in wild-type but not S310F, contrasting with our *in vitro* finding that T-DXd co-treatment had little effect. We experimentally tested the docking prediction that patritumab engages HER3 via ECD1 by generating HA-tagged HER3 truncations lacking individual extracellular domains and expressing them in UACC3133 and BCK4 (**Figure 6f-g**). Using P-DXd for immunoprecipitation, full-length HER3, ΔECD3, and ΔECD4 co-precipitated efficiently, whereas ΔECD1 and ΔECD2 failed to bind (**Figure 6h**). We asked which domains are necessary for the P-DXd-driven increase in HER2/HER3 association. P-DXd robustly increased HER2/HER3 interaction with full-length HER3 and with ΔECD3 or ΔECD4, but this effect was lost with ΔECD1 and ΔECD2 (**Figure 6i**). These data indicate that both ECD1 and ECD2 are required for P-DXd binding and P-DXd-induced HER2/HER3 association.

Having shown that P-DXd binds HER3 ECD1 and promotes HER2/HER3 complex formation, we asked how this interaction affects T-DXd-induced HER2 ubiquitination. In UACC3133 and BCK4, T-DXd increased HER2 ubiquitination, co-treatment with P-DXd further amplified it, and P-DXd alone caused a modest rise (**Figure 6j**). These data indicate that P-DXd engagement of HER3 favors HER2 ubiquitin tagging, thereby potentiating T-DXd-mediated HER2 turnover. We addressed whether this affects T-DXd trafficking. In endogenous and isogenic models, co-treatment with P-DXd accelerated T-DXd internalization in a dose-dependent manner across mutants and wild-types, consistent with the ubiquitination results (**Figure 6k-l**). We determined whether both ADCs are co-internalized in HER2-mutant ILC through shared endocytic compartments (**Figure 6m**). After simultaneous exposure to T-DXd and P-DXd, HER2-mutant cells showed strong spatial overlap of the two ADC signals.

To determine whether the observed effects were specific to P-DXd or intrinsic to its antibody component, we compared patritumab with P-DXd in HER2-mutant ILC models. Patritumab blocked HRG-driven signaling and increased HER2/HER3 complex formation, mirroring P-DXd (**Extended Data 6a**). Both agents similarly enhanced HER2 ubiquitination, indicating that the antibody alone can induce receptor conformations favoring ubiquitin tagging and turnover (**Extended Data 6b**). We assessed whether patritumab modulates T-DXd trafficking. Co-treatment with patritumab and T-DXd further increased HER2 ubiquitination compared with either agent alone (**Extended Data 6c**), and patritumab accelerated T-DXd internalization in a dose-dependent manner in both mutant and wild-type models (**Extended Data 6d-e**). Functionally, T-DXd plus patritumab produced moderate synergy in UACC3133 and IPM-BO-030 and mainly additive effects in BCK4 (**Extended Data 6f**). The synergy was less pronounced than T-DXd/P-DXd combination, suggesting the ADC form provides added benefit through its cytotoxic payload.

To evaluate *in vivo* efficacy, we established orthotopic xenografts with UACC3133 and BCK4. Mice received vehicle, T-DXd, P-DXd, or the combination, a regimen within standard human exposures. In UACC3133, both single agents significantly suppressed growth during dosing (**Figure 7a-b**). The combination produced only a modest, non-significant additional reduction on treatment, but clear differences emerged after dosing stopped. Tumors in the combination arm remained durably controlled, whereas P-DXd-treated tumors resumed growth and a subset in the T-DXd arm regrew near the end of the 100-day follow-up (**Figure 7d**). All active arms extended survival versus vehicle, with T-DXd and the combination outperforming P-DXd monotherapy (**Figure 7c**). BCK4 showed a similar pattern, with robust on-treatment suppression by each single agent and comparatively greater P-DXd activity than in UACC3133, consistent with higher *in vitro* sensitivity (**Figure 7e-f**). After cessation, the combination maintained control, whereas P-DXd tumors gradually regrew and a tumor in the T-DXd arm also regrew (**Figure 7h**). Survival improved across all active arms (**Figure 7g**). Across all arms, weights remained stable and comparable to vehicle throughout. No animals exhibited distress, ruffled fur, reduced activity, or other behavioral abnormalities, indicating that neither single-agent nor combination therapy produced overt systemic toxicity (**Supplementary Figure 6a-b**). Because the *in vivo* studies did not include a high-dose T-DXd arm matching the total ADC dose of the combination, we performed a complementary *in vitro* experiment (**Extended Data 7a-c**). Across endogenous HER2-mutant models, dual-ADC treatment more completely suppressed pHER2/pHER3/pAKT/pERK, produced higher pH2AX, and caused greater viability loss than T-DXd alone at the same total concentration. These results indicate that the advantage of T-DXd plus P-DXd reflects true biological synergy rather than simple dose additivity.

**Figure 7.**
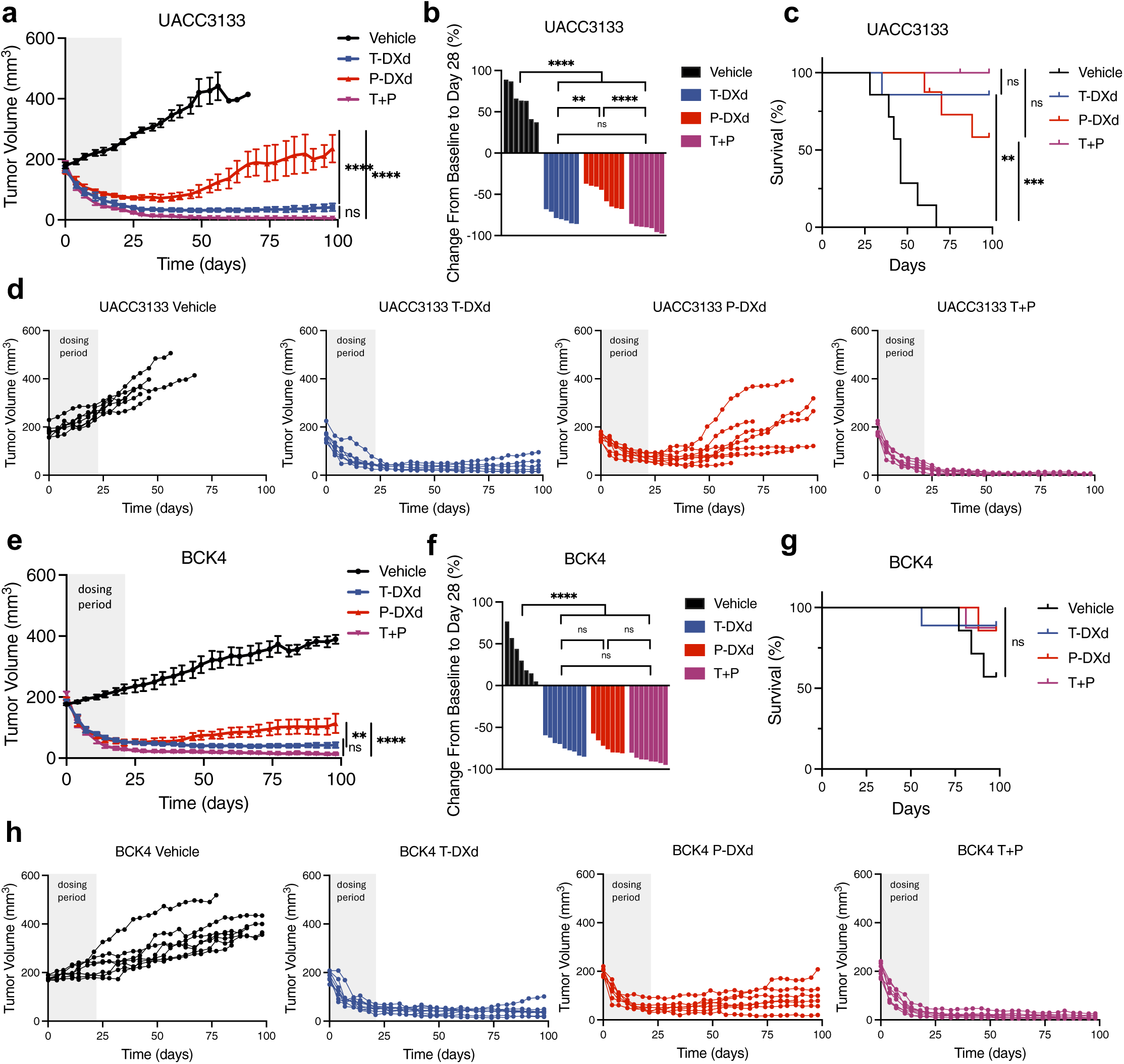
Combination of T-DXd and P-DXd suppresses tumor growth in HER2-mutant ILC xenografts. (a) Tumor growth curves of UACC3133 xenografts treated with vehicle, T-DXd (2.5 mg/kg), P-DXd (2.5 mg/kg), or the combination (T + P; 5 mg/kg total). Error bars represent mean ± SEM. (b) Individual tumor growth curves for each treatment group of UACC3133 xenografts. (c) Percent tumor volume change from baseline to day 28 across treatment arms for UACC3133 xenografts. (d) Kaplan-Meier survival analysis of UACC3133 xenografts comparing all treatment groups to vehicle control. (e) Tumor growth curves of BCK4 xenografts treated with vehicle, T-DXd (2.5 mg/kg), P-DXd (2.5 mg/kg), or the combination (T + P; 5 mg/kg total). Error bars represent mean ± SEM. (f) Individual tumor growth curves for each treatment group of BCK4 xenografts. (g) Percent tumor volume change from baseline to day 28 across treatment arms for BCK4 xenografts. (h) Kaplan-Meier survival analysis of BCK4 xenografts comparing all treatment groups to vehicle control. Statistical significance was determined using one-way or two-way ANOVA; *p* < 0.05.

To assess the breadth and robustness of T-DXd plus P-DXd, we expanded testing to additional PDOs with endogenous hotspot HER2 mutations and models with T-DXd resistance (**Supplementary Table 1, Supplementary Figure 7**). In two NST PDOs, IPM-BO-124 with HER2 amplification plus S310F and IPM-BO-222 with HER2 amplification plus L755S, both responded to T-DXd, but with greater potency in IPM-BO-222 (**Extended Data 8a**). P-DXd alone showed minimal activity. The combination produced synergistic growth inhibition that exceeded either monotherapy and yielded greater suppression of HER2/HER3/AKT/ERK phosphorylation, and greater pH2AX accumulation than an equal total dose of T-DXd alone (**Extended Data 8b-c**). To model clinically relevant T-DXd resistance, we used PDOs from patients who progressed on T-DXd, including IPM-BO-317B and IPM-BO-318C with intrinsic resistance and IPM-BO-315D with acquired resistance (**Supplementary Table 1, Supplementary Figure 7**). We confirmed that resistance persisted *ex vivo* by benchmarking these PDOs against a broader panel with previously unknown sensitivity to T-DXd and P-DXd (**Extended Data 9a**). IC50 values separated the cohort into T-DXd-sensitive and T-DXd-resistant groups, and all three autopsy-derived models clustered in the resistant set. P-DXd monotherapy showed heterogeneous activity across PDOs, indicating variable HER3 dependence.

We profiled molecular correlates of T-DXd resistance across PDOs (**Extended Data 9b-c**). HER2 mRNA and protein were markedly reduced in resistant models, consistent with limited target availability. HER3 expression remained relatively stable, aligning with the more uniform P-DXd activity. Despite lower total HER2, several resistant PDOs retained pHER2 (and pHER3) at levels comparable to sensitive models, indicating maintained HER2/HER3 signaling in a subset. EGFR and MRP1 were elevated in resistant IPM-BO-317B and IPM-BO-292B. These two PDOs also showed attenuated HER2/HER3 activity, suggesting an EGFR/MRP1-driven bypass. We tested T-DXd/P-DXd combination in these three T-DXd-resistant PDOs (**Extended Data 9d**). The combination showed robust synergy in all models, with more complete suppression of pHER2/pHER3/pAKT/pERK and stronger pH2AX induction than an equal total dose of T-DXd alone (**Extended Data 9e**).

To examine how acquired T-DXd resistance evolves in HER2-wild-type versus HER2-mutant settings, we generated T-DXd resistant (TR) derivatives from IPM-BO-053 wild type, S310F, and V777L under chronic T-DXd exposure (**Extended Data 10a-b**) Immunoblotting showed variant-specific remodeling (**Extended Data 10c**). HER2 increased in wild-type and S310F HER2 mutant but decreased in V777L HER2 mutant, while pHER2 rose in all, indicating sustained activation despite divergent expression. HER3 decreased in wild-type and S310F HER2 mutant and increased in V777L HER2 mutant; pHER3 was elevated across derivatives, suggesting convergent HER3 activation as a compensatory axis. Total and phosphorylated EGFR declined in all resistant lines, arguing against an EGFR bypass. MRP1 was uniformly upregulated. qRT-PCR showed partial mismatch with protein levels (**Extended Data 10d**). HER2 mRNA rose modestly in wild-type and S310F TR lines and was unchanged in V777L TR. HER3 mRNA increased only in V777L TR. MRP1 transcripts were uniformly elevated. Combined with protein data, this supports mutation-specific resistance programs, with wild type and S310F aligning and V777L diverging, and with recurrent nodes of HER2/HER3 activation and MRP1 upregulation across genotypes. Across the two isogenic T-DXd-resistant HER2-mutant PDOs, T-DXd plus P-DXd remained strongly synergistic, with more complete suppression of pHER2/pHER3/pAKT/pERK and greater induction of pH2AX than either monotherapy or dose-matched T-DXd (**Extended Data 10e-f**).

## Discussion

HER2-mutant ILC remains an unmet need despite clear genomic enrichment and poor outcomes. Here we show that T-DXd is highly active in this subtype and that efficacy is driven not by target abundance but by mutation-conditioned receptor fate. Activating mutations shift HER2 from a relatively stable, recycling receptor to one that is preferentially ubiquitinated, internalized, and lysosomally degraded after ligand or T-DXd engagement, thereby boosting T-DXd delivery and creating a lever for combinations. The inclusion of both ER⁺/HER2-mutant and ER⁻/HER2-mutant ILC models expands the translational scope of this work and enables future testing of endocrine/T-DXd co-therapy strategies in ER+ disease.

Traditional HER2 therapy emphasizes how much HER2 is on the surface and how dependent the tumor is on its kinase activity [34, 35]. In HER2-mutant ILC, those measures are not sufficient, and T-DXd therapeutic benefit is additionally influenced by post-binding receptor dynamics. Modest HER2 destabilization with an irreversible HER2 TKI or HSP90 inhibition augments T-DXd engagement without exhausting the target. The lack of striking synergy in short-term assays and the clear gains in long-term survival assays are not a contradiction; they reflect ILC biology and the time it takes DNA damage to accrue in slowly cycling cells [36, 37]. A practical strategy is sub-cytotoxic priming to increase ubiquitin tagging and endocytic uptake of T-DXd. Consistent with this concept, T-DXd plus neratinib has demonstrated antitumor activity with manageable toxicity in advanced HER2-altered solid tumors [38], and ganetespib may offer a better- tolerated alternative based on breast cancer trial experience [39–42]. Trial designs should test low-intensity priming given concurrently or as a brief pre-treatment run-in, rather than dose-escalating toward complete pathway quiescence.

Our study demonstrates preferential HER3 activation in HER2-mutant ILC. Functionally, HER2-mutant ILC cells show increased HER2/HER3 association along with enhanced HER3 ubiquitination and internalization. HER3 knockdown impairs growth to a similar extent as HER2 loss, and combined depletion produces the strongest effect, supporting co-dependence of HER2 and HER3. This activated HER3 state is therapeutically actionable with P-DXd, which shows strong synergy with T-DXd *in vitro* and, in combination, prevents recurrence *in vivo*. The surprising mechanistic observation is that P-DXd increases HER2/HER3 complex formation and promotes HER2 ubiquitination and internalization while suppressing signaling. Epitope mapping points to a dual requirement for HER3 ECD1 and ECD2, consistent with a drug-organized dimer that is signaling-inert yet trafficking-competent. This mirrors prior reports of drug-induced HER2 heterodimerization with HER3 or EGFR, and provides a structural rationale for synergy that exceeds DXd dose additivity [43–45]. Rather than blocking dimerization, HER3 engagement choreographs a conformation optimized for co-internalization and lysosomal processing of both ADCs. Early clinical experience supports dual HER2/HER3 targeting, with durable responses to trastuzumab plus pertuzumab reported in patients harboring concurrent HER2 and HER3 mutations, including an ILC case [46].

Patient-derived organoids with clinical resistance show three recurring themes: target loss limits binding, efflux limits payload retention, and signaling rewires toward residual HER2/HER3 activity or, in some contexts, toward EGFR. The dual-ADC strategy addresses at least two of these. By engaging HER3, it may help re-establish a trafficking-competent receptor complex even when HER2 expression is diminished, while coordinated co-internalization could increase intracellular payload delivery sufficiently to partially overcome efflux-mediated loss. Beyond dual targeting alone, payload diversification may provide an additional means of circumventing resistance, as supported by the feasible combination of sacituzumab govitecan and enfortumab vedotin in metastatic urothelial carcinoma [47]. If MRP1 is a recurrent efflux mechanism, then alternating, sequencing, or combining payloads with distinct transporter liabilities could prolong benefit while minimizing overlapping toxicities. At the same time, combinations of distinct ADCs carrying a shared payload have also shown promise as a resistance-overcoming strategy. Recent work demonstrated that T-DXd resistance can arise through HER2 loss or trastuzumab-binding interface mutations that impair ADC binding and internalization, and that combining low-dose T-DXd with a TROP2-directed ADC restored more uniform intratumoral DXd delivery and overcame HER2-loss-mediated resistance [48]. Ultimately, an especially attractive design may be a dual-payload HER2×HER3 bispecific ADC that stabilizes a signaling-inert, rapidly internalized heterodimer while efficiently delivering cytotoxic cargo. In this setting, rational selection of payloads less susceptible to MRP1-mediated export, such as MMAF, may further extend benefit in efflux-high tumors [49].

Three trial concepts follow directly. First, a single-arm signal-seeking study of T-DXd in HER2-mutant ILC with built-in pharmacodynamics to validate biomarkers. Second, a randomized trial that tests T-DXd plus P-DXd versus T-DXd alone, using alternating or staggered schedules and conservative dosing to manage interstitial lung disease risk. Third, a salvage cohort enriched for clinical T-DXd resistance, where the question is whether the combination restores sensitivity by re-establishing a trafficking-competent state.

Key limitations remain. The Caris analysis is retrospective and subject to selection bias and unmeasured confounding factors. Survival is unadjusted for key covariates, and T-DXd timing/line varied, risking confounding by indication and limiting causal inference. *In vitro* model systems capture receptor-intrinsic mechanisms but not the full complexity of tumor microenvironment, stromal trafficking cues, or immune engagement that can influence ADC performance. Prospective work should include spatial proteomics and live-imaging in more physiologic *in vivo* systems to quantify receptor fate and payload distribution. Finally, although our *in vivo* studies support tolerability at conservative doses, clinical translation of dual-DXd regimens will likely hinge on dose optimization and schedule refinement. Accordingly, lower-dose combination and sequential dosing paradigms are being actively evaluated to mitigate overlapping toxicities while preserving efficacy.

In summary, HER2 mutations in ILC shift HER2 toward ubiquitination, internalization, and lysosomal routing, making T-DXd highly effective. Low-dose priming with neratinib or ganetespib further increases HER2 ubiquitin tagging, yielding durable long-term control. Concurrent HER3 engagement with P-DXd stabilizes a signaling-inert yet trafficking-competent HER2/HER3 complex, enabling cooperative dual-ADC uptake. Together, these insights support a biology-guided strategy to enhance T-DXd response and delay resistance.

## Materials and Methods

### Clinical data sets

Breast cancer tumors that went through comprehensive molecular profiling tests at Caris Life Sciences (Phoenix, AZ) were included as the clinical data set. Real-world clinical data were obtained from insurance claims. Caris CODEai advanced breast cancers with centralized profiling, curated clinicopathologic data, and survival follow-up were analyzed retrospectively. Histology was classified as ILC or NST; ER+ was defined as ≥ 1 % by IHC; HER2 non-amplified status was determined by copy number. Somatic HER2 variants from NGS were COSMIC-aligned and Caris-curated; synonymous variants and VUS were excluded, retaining pathogenic/likely pathogenic calls. Within ER+/HER2-nonamplified tumors, HER2-mutation prevalence between ILC and NST and overall survival within each histology were evaluated. T-DXd outcomes were assessed as survival from treatment start to last contact within each histology, comparing HER2-mutant versus wild type.

### Cell lines and patient-derived organoids

MDA-MB-134-VI (MM134) and UACC3133 were from ATCC; SUM44PE from BioIVT; BCK4 were obtained under MTA from Dr. Britta Jacobsen (University of Colorado Anschutz) and can not be distributed further; IPH926 from Dr. Matthias Christgen (Hannover Medical School); and HCC2185 from Dr. Adi Gazdar (UT Southwestern). Cells were cultured with 10% FBS in the following media: MM134 in 1:1 DMEM:L-15, UACC3133 in DMEM, IPH926 and HCC2185 in RPMI, BCK4 in MEM with non-essential amino acids and insulin; SUM44PE in DMEM/F12 with 2 % charcoal-stripped serum plus supplements [54, 116]. Patient-derived organoids were obtained from the Institute for Precision Medicine (University of Pittsburgh) and maintained in breast organoid medium [117]. All cell lines and PDOs were Mycoplasma-negative and cell lines were authenticated by STR profiling at the Genomics Core of the University of Arizona.

### CRISPR/Cas9-edited HER2 mutation knock-in models

HER2 mutations S310F and V777L were introduced by CRISPR/Cas9 prime editing. pegRNAs and nicking sgRNAs were designed to install the desired nucleotide changes and minimize bystander edits using an online tool (PrimeDesign). Cells or single-cell PDO suspensions were transfected with PE2 plasmid plus pegRNA and nicking sgRNA using FuGENE HD followed by spinoculation. Forty-eight hours post-transfection, GFP+ cells were single-cell sorted into 96-well plates. Clones were expanded, genomic DNA was PCR-amplified, and Sanger sequencing verified the edit. Matched wild-type isogenic controls were generated using non-editing pegRNAs and the same selection pipeline. Multiple independent clones per genotype were pooled to avoid clonal drift

### siRNA knockdown

Cells were transfected with *ERBB2* or *ERBB3* siRNAs (Dharmacon) using Lipofectamine RNAiMAX per manufacturer instructions, with a non-targeting siRNA as control. Cells were collected 72 h later for immunoblotting and viability assays.

### Compounds

Neratinib, afatinib, tucatinib, poziotinib, and ganetespib, dyngo-4a, cycloheximide, bafilomycin A1 and MG132 were dissolved in DMSO. Trastuzumab deruxtecan, patritumab deruxtecan, trastuzumab, patritumab, pertuzumab, LJM716, and zenocutuzumab were prepared in PBS. Drugs were purchased from MedChemExpress.

### Drug response assay

Cells were seeded at 1,000 per well in 96-well plates and treated 24 h later with drug or vehicle for 7 days. Viability was quantified with PrestoBlue HS and normalized to vehicle. For PDO assays, organoids were dissociated to single cells and plated at 1,000 cells in 5 µL Cultrex per well in 96-well round-bottom plates. Drugs or vehicle were added 3 days after plating, and viability was measured on day 10 post-treatment with CellTiter-Glo 3D. Dose-response curves were fit by nonlinear regression in GraphPad to derive IC50 values. For short-term synergy, drugs were combined across serial dilutions and synergy was quantified with the HSA model. For long-term synergistic effects, clonogenic assays were done by plating 1,000 cells per well in 12-well plates, treating for 21-28 days, fixing in methanol, staining with 0. 5% crystal violet, and dissolving the dye for absorbance-based colony quantification.

### Immunoblotting

Cells or PDOs were lysed in RIPA buffer with protease/phosphatase inhibitors. Equal protein was resolved by SDS-PAGE and transferred to PVDF, and probed with primary antibodies overnight at 4 ° C. After washes, membranes were incubated with fluorescent or HRP-conjugated secondary antibodies for 1 hour at room temperature. Fluorescent signals were imaged on a LI-COR Odyssey CLx; chemiluminescence was captured on a Bio-Rad ChemiDoc. Band intensities were quantified with the instrument software or ImageJ.

### Co-immunoprecipitation

For HER2 /HER3 interaction assays, cells or PDOs were serum-starved then stimulated with HRG or drugs. For ubiquitination assays, samples were pretreated with bafilomycin A1 or vehicle and then exposed to HRG or drugs. Cells were lysed in IP buffer with protease/phosphatase inhibitors and quantified. Immunoprecipitation used Dynabeads Protein G coupled to HER2 or HER3 antibody for co-IP and ubiquitination studies, or to P-DXd for HER3 epitope binding, and were then incubated with clarified lysates.

Immune complexes were washed, eluted, resolved by SDS-PAGE, transferred to PVDF, and immunoblotted for HER2, HER3, ubiquitin, rabbit IgG and human IgG. Input lysates were run in parallel, and band intensities were quantified by ImageJ.

### Proximity ligation assay

HER2/HER3 proximity was measured with the NaveniFlex Cell MR Atto647N per manufacturer instructions. Cells were fixed in 4 % PFA, permeabilized, blocked, and incubated with primary antibodies raised in different species to HER2 and HER3. After ligation and amplification, nuclei were counterstained with DAPI. Foci per nucleus were quantified from ≥6 fields per condition.

### ADC internalization

ADCs were pHrodo-labeled per kit instructions. For internalization assays, cells were seeded in clear-bottom black 96-well plates and incubated at 37°C with 1ug/ml pHrodo-T-DXd or pHrodo-P-DXd. Time-lapse images were collected on an Incucyte; where indicated, cells were pretreated with dyngo-4a. Mean fluorescence was normalized to confluence and quantified over time. For lysosomal routing, cells were co-incubated with LysoBrite Green and colocalization with pHrodo-ADC signal was quantified in ImageJ using Pearson’s coefficients.

### qRT-PCR

Total RNA was extracted with RNeasy. cDNA was synthesized using PrimeScript RT Master Mix. qPCR was done with SsoAdvanced Universal SYBR, and relative expression was calculated by 2 ^ -ΔΔCt normalized to RPLP0.

### Molecular docking

HER2/HER3 complex structures 7MN5 and 7MN8 were downloaded from RCSB PDB; HRG segments were removed in PyMOL. As no experimental crystal structure exists for patritumab, the Fab structure was predicted using AlphaFold Server (ipTM = 0.93, ranking score = 0.89) from the heavy chain (449 aa) and light chain (219 aa) sequences; chain IDs were reassigned to antibody convention (H/L). Complexes were relaxed with Rosetta FastRelax, docked via ClusPro, and the top pose was solvated/ionized in CHARMM-GUI for GROMACS. MD (GROMACS 2023.2) followed the CHARMM-GUI workflow: minimization, staged equilibration, and 200-ns production. Trajectories were concatenated, made whole, de-imaged, and protein-fitted (trjcat/trjconv). Binding energies were computed with gmx_MMPBSA (PB model, 0.15M, 310.15K), excluding glycans unless noted; chain IDs were standardized and index groups defined for HER2, HER3, and antibodies. For antibody systems, ΔG was calculated separately for patritumab-HER3 and trastuzumab-HER2. Structures were visualized in VMD 1.9.3.

### Lentiviral transduction

pCDH-CMV-EF1A-EGFP-T2A-PURO vectors encoding full-length HER3 or ΔECD1–ΔECD4 were packaged in HEK293T with pPAX2 and pMD2.G using FuGENE 6. Filtered supernatants (48-72 h) transduced target cells with 8 µg/mL polybrene. Stable pools were selected in 1 µg/mL puromycin; expression was confirmed by immunoblot.

### Generation of T-DXd-resistant PDOs

T-DXd-resistant PDOs were generated from IPM-BO-053 models harboring wild-type HER2, HER2 S310F, or HER2 V777L. Parental PDOs were maintained under standard organoid culture conditions and chronically exposed to escalating doses of T-DXd for approximately 6 months. Surviving organoids were allowed to recover and expand during treatment, enriching for resistant populations. At the end of the selection period, surviving organoids were expanded in culture and established as T-DXd-resistant PDO derivatives for downstream functional and molecular analyses. Parental vehicle-treated control PDOs were cultured in parallel for the same duration.

### *In vivo* studies

All procedures were IACUC-approved. Female NSG mice (8 weeks) received orthotopic injections of 5×10⁶ UACC3133 or BCK4 cells into the right mammary fat pad. A 0.36-mg 17β-estradiol (E2) pellet was implanted; E2 (8 μg/mL) was later supplemented in drinking water. When tumors reached 150-200 mm³, mice were randomized and received vehicle, T-DXd (2.5 mg/kg), P-DXd (2.5 mg/kg), or the combination (2.5+2.5 mg/kg) once weekly ×4 via tail vein. Tumors were measured twice weekly; volume = (L×W²)/2. Body weight/clinical status were monitored; survival was followed to 100 days.

### Statistical analysis

Unless noted, experiments were performed in biological triplicate with technical replicates. Data are mean ± SD/SEM. IC50 values were fit by nonlinear regression; curves compared by global F-test. Groups were analyzed by two-tailed t-tests or one/two-way ANOVA with post hocs. Synergy used HSA (SynergyFinder), with scores >10 deemed synergistic. Significance was set at P<0.05.

## Acknowledgements

We thank all patients who participated in the Hope for OTHERS program and donated tissue for this study. We thank the University of Pittsburgh Organoid Research Core Facility (RRID: SCR_025698) for generating the PDOs, the University of Pittsburgh Animal Facility for assistance with animal studies, and the University of Pittsburgh Center for Research Computing Core (RRID: SCR_022735) which is supported by NIH award S10OD028483 (to A.V.L.).

## Contributions

J.L. designed and performed the main experiments. Y.C. assisted with mouse studies and bioinformatics analyses. A.C. assisted with molecular modeling and binding energy analyses, and R.J. provided clinical data. D.B. and H.W. generated PDOs, and D.T. assisted with experiments. R.M.J., G.N., J.H., J.F., and M.B. provided input and suggestions. J.L. analyzed the data, interpreted the results, and wrote the manuscript. All authors edited the manuscript. A.V.L. and S.O. supervised and oversaw the study.

## Funding

This project was supported by the National Institutes of Health under awards 5F30CA278319-03 (to J.L.) and R01CA252378 (to S.O. and A.V.L.), by the Breast Cancer Research Foundation (to S.O.), and in part by Puma Biotechnology.

## Competing interests

J.L. has received funding from Puma Biotechnology. R.J. is an employee of Caris Life Sciences. G.N. has received speaker honoraria from AstraZeneca and Novartis. J.F. has received institutional funding from the Pennsylvania Breast Cancer Coalition and is supported by the National Center for Advancing Translational Sciences of the National Institutes of Health under Award Number KL2TR001856. The content is solely the responsibility of the authors and does not necessarily represent the official views of the National Institutes of Health. J.F. has also received consulting fees and/or honoraria from OncLive, Biotheranostics, Inc., a Hologic company, Genentech, Daiichi Sankyo, Gilead, AstraZeneca, Novartis, and Caris Life Sciences.

## Figure Legends

**Extended Data 1.**
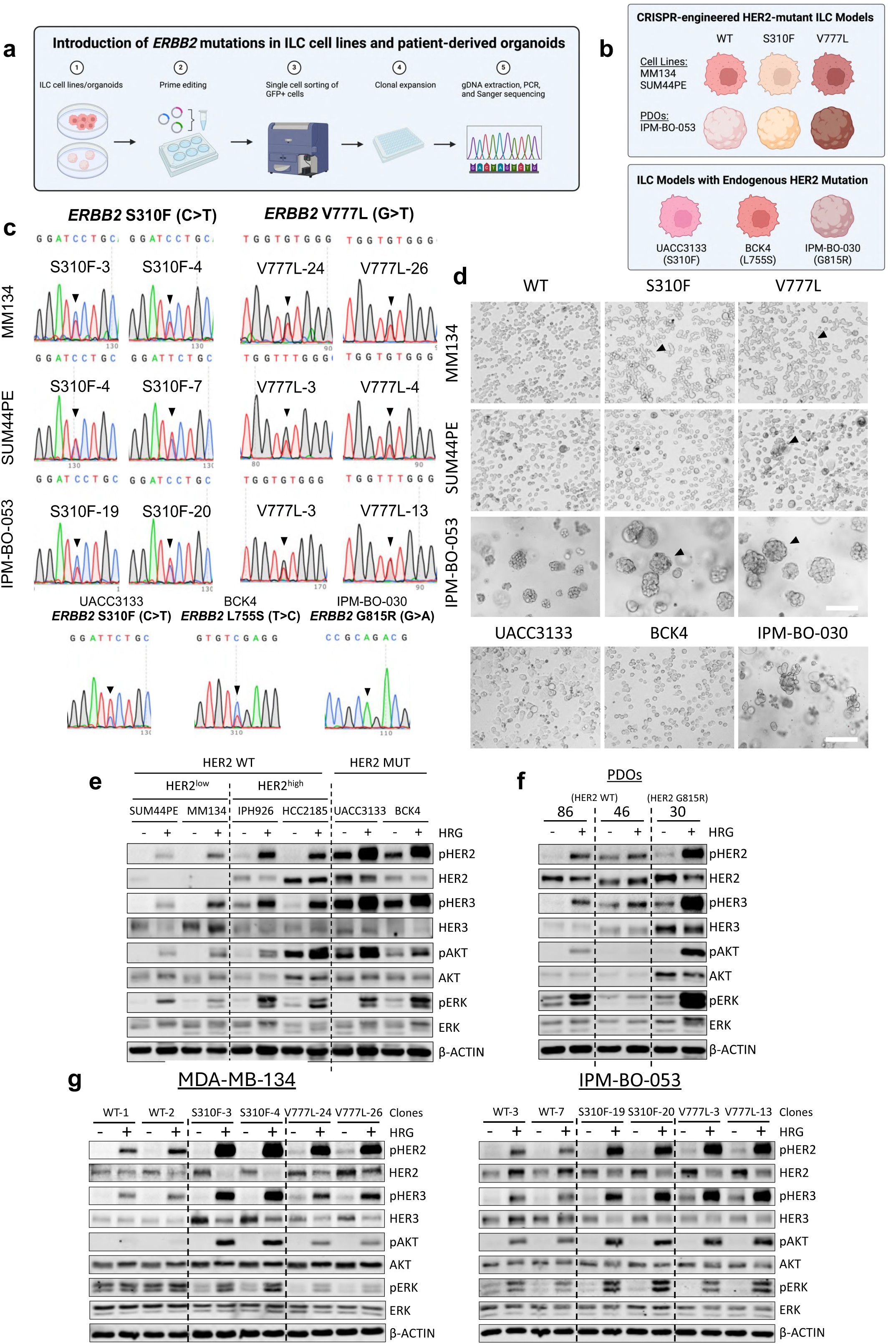
ILC cell line and patient-derived organoid (PDO) models with CRISPR knock-in or endogenous HER2 mutations show hyperactive HER2/HER3 and downstream signaling. (a) Workflow schematic of CRISPR/Cas9 prime editing used to introduce activating HER2 mutations at the endogenous locus. (b) Summary of HER2-mutant ILC models generated or identified. (c) Representative Sanger sequencing traces confirming targeted HER2 substitutions or endogenous mutations. (d) Representative bright-field images showing morphological differences of isogenic HER2-mutant versus wild-type cells and PDOs. Scale bar, 300 µm. (e) Immunoblot analysis of HER2 and downstream signaling under serum-starved and HRG-stimulated conditions (1 nM, 10 min) in endogenous HER2-wild-type (low- and high-HER2-expressing) and HER2-mutant ILC cell lines. (f) Immunoblot analysis of HER2 and downstream signaling under serum-starved and HRG-stimulated conditions (1 nM, 30 min) in endogenous HER2-wild-type PDOs (IPM-BO-086 and IPM-BO-046) and HER2-mutant PDO (IPM-BO-030). (g) Immunoblot analysis of HER2 and downstream signaling under serum-starved and HRG-stimulated conditions (1 nM, 10 min) in isogenic MM134 and IPM-BO-053 wild-type and mutant clones

**Extended Data 2.**
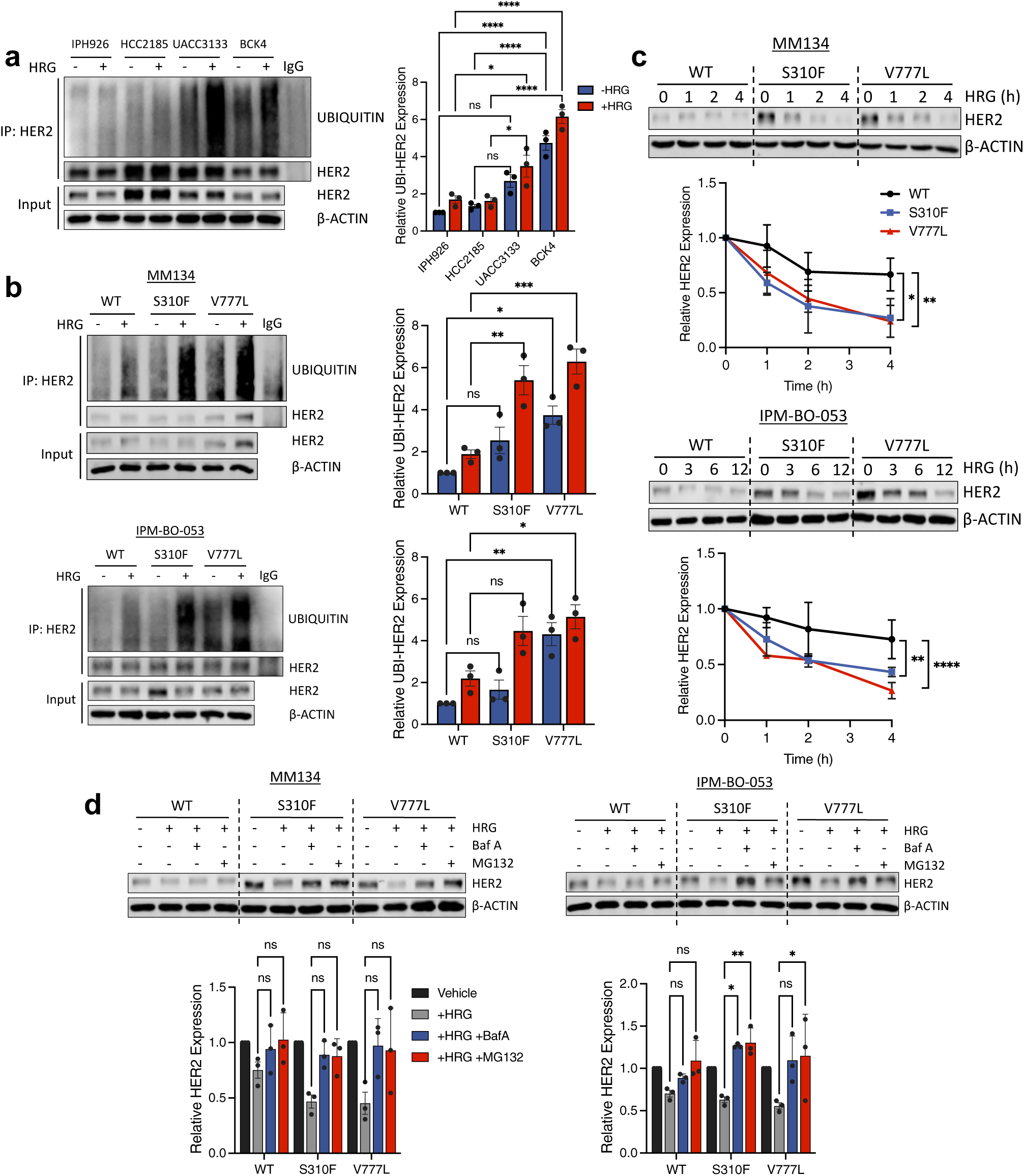
Activating HER2 mutations in ILC enhance HRG-induced ubiquitination and lysosomal degradation of HER2. (a) Immunoblot analysis of HER2 ubiquitination in serum-starved endogenous HER2-mutant (UACC3133, BCK4) and HER2-wild-type ILC cell lines (IPH926, HCC2185) pretreated with bafilomycin A1 (Baf A1; 100 nM, 2 h) followed by vehicle or HRG (1 nM, 4 h), with quantification normalized to vehicle-treated IPH926. Error bars represent mean ± SEM. (b) Immunoblot analysis of HER2 ubiquitination in serum-starved isogenic MM134 and IPM-BO-053 models pretreated with Baf A1 (100 nM, 2 h) followed by vehicle or HRG (1 nM, 4 h), with quantification normalized to vehicle-treated wild-type controls. Error bars represent mean ± SEM. (c) Cycloheximide chase assay comparing HER2 degradation kinetics in serum-starved isogenic models following HRG (1 nM) stimulation, with HER2 expression normalized to baseline (t = 0). Error bars represent mean ± SEM. (d) Pharmacologic stabilization assays comparing serum-starved isogenic models pretreated with Baf A1 (100 nM, 2 h) or MG132 (20 µM, 2 h), followed by HRG (1 nM, 4 h), with HER2 expression normalized to vehicle. Error bars represent mean ± SEM. Representative results are shown from three independent experiments (n = 3). Statistical significance was determined using two-way ANOVA; *p < 0.05.

**Extended Data 3.**
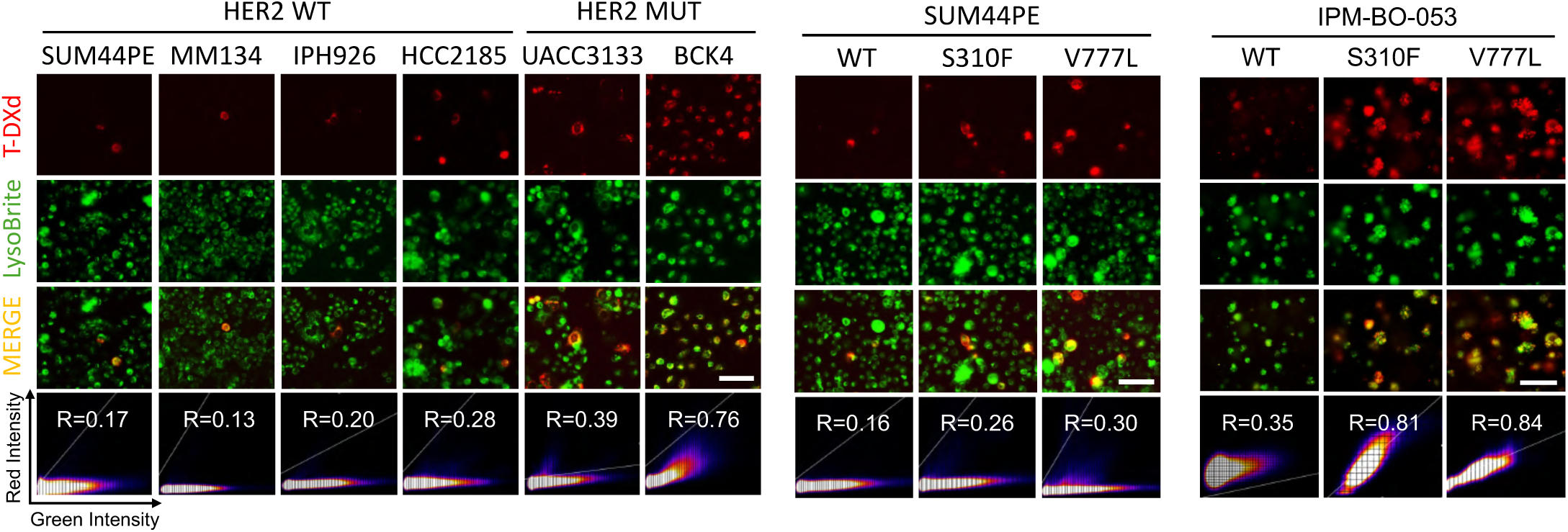
T-DXd traffics to lysosomes. Live-cell fluorescence imaging of pHrodo-T-DXd (1 µg/mL; red) and LysoBrite™ Green-labeled lysosomes (10 nM) in endogenous and isogenic HER2-mutant models after 48 hours. Fluorescent signal co-localization was quantified using Pearson’s correlation coefficient. Representative results are shown from three independent experiments (n = 3). Scale bar, 300 µm.

**Extended Data 4.**
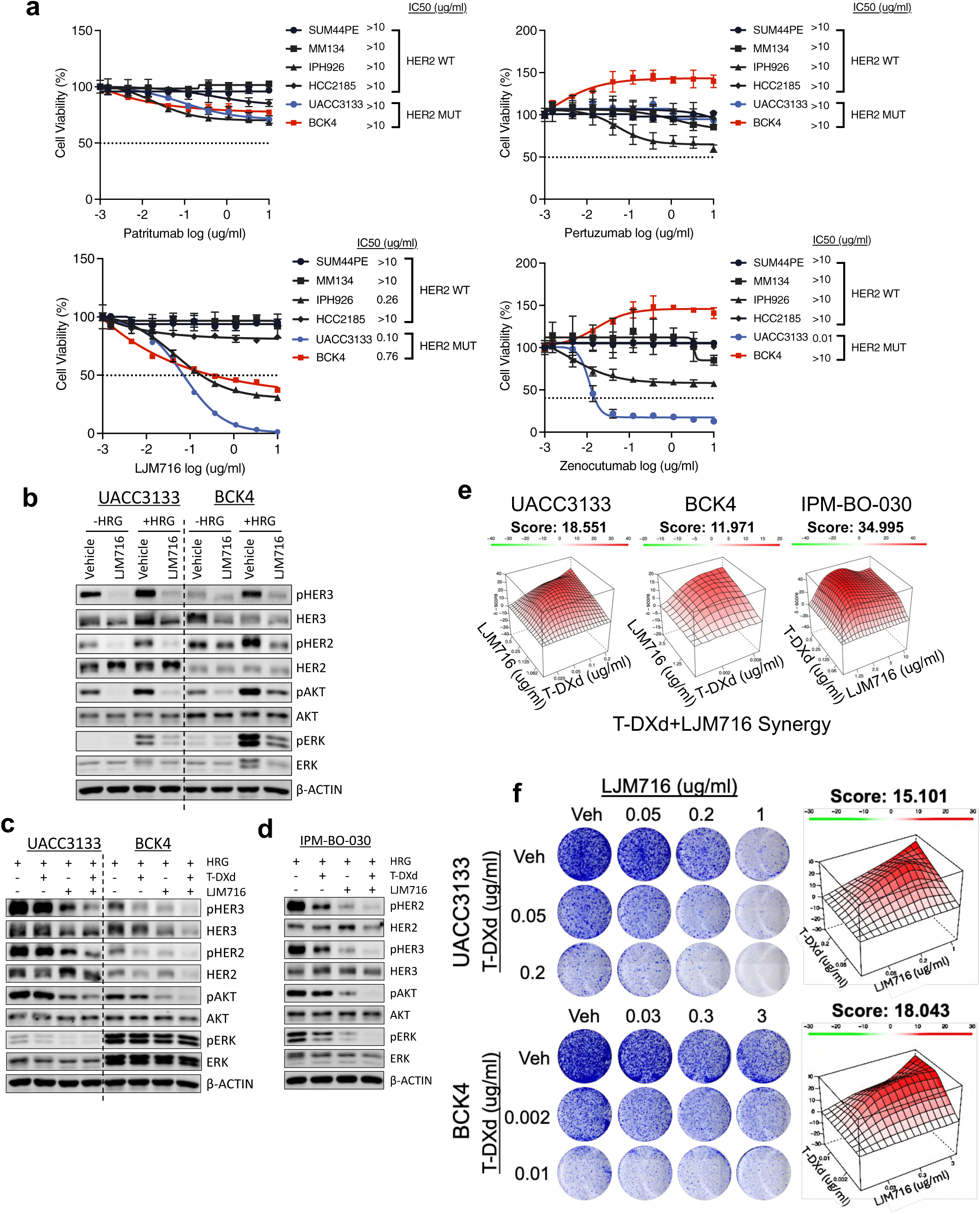
Sensitivity to additional HER3-targeting agents (patritumab, pertuzumab, LJM716, zenocutuzumab). (a) Dose-response curves comparing sensitivity of endogenous HER2-mutant (UACC3133, BCK4) and HER2-wild-type (SUM44PE, MM134, IPH926, HCC2185) ILC cell lines to HER3-directed agents (LJM716, patritumab, pertuzumab, zenocutuzumab). Error bars represent mean ± SD. (b) Immunoblot analysis of HER2/HER3 and downstream signaling inhibition in serum-starved UACC3133 and BCK4 pretreated with LJM716 (10 µg/mL, 4 h) followed by vehicle or HRG (1 nM, 10 min). (c-d) Immunoblot analysis of HER2/HER3 and downstream signaling inhibition in serum-starved UACC3133 and BCK4 (c), IPM-BO-030 (d), pretreated with T-DXd (1 µg/mL, 4 h), LJM716 (1 µg/mL, 4 h), or the combination, followed by vehicle or HRG (1 nM, 10 min or 30 min). (e) Short-term (7-day) proliferation analysis of T-DXd, LJM716, and the combination in UACC3133, BCK4, and IPM-BO-030. (f) Long-term (3-4 week) clonogenic survival analysis of T-DXd, LJM716, and the combination in UACC3133 and BCK4. Representative results are shown from three independent experiments (n = 3). Synergy scores were calculated using the HSA model; values > 10 indicate synergy.

**Extended Data 5.**
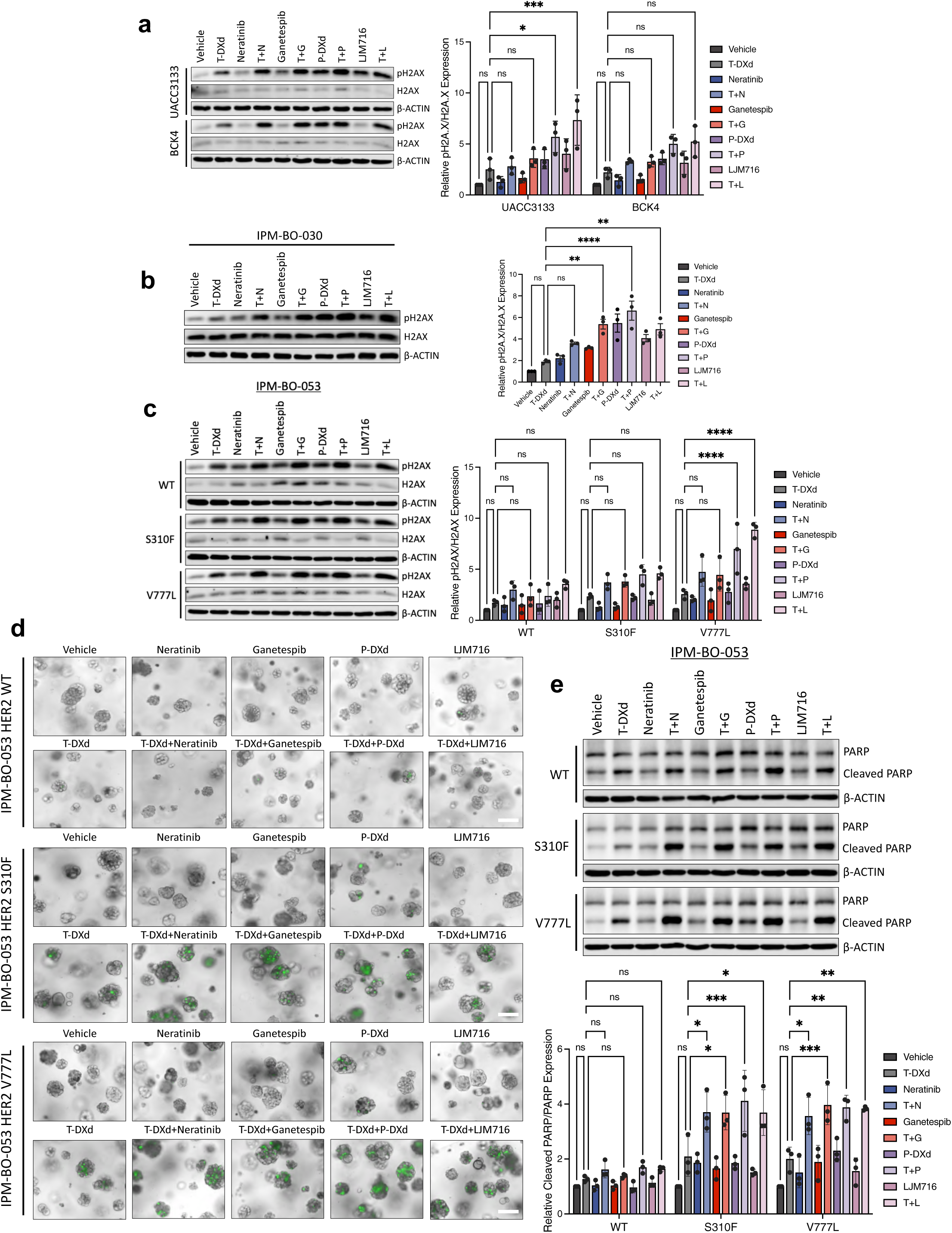
T-DXd-based combination treatments enhance DNA damage and apoptosis in HER2-mutant ILC. (a-c) Immunoblot analysis of pH2AX and H2AX following 72-hour single-agent or combination treatments in UACC3133 and BCK4 (a), IPM-BO-030 (b), and IPM-BO-053 isogenic models (c), with quantification of pH2AX/H2AX ratios normalized to vehicle-treated controls. Error bars represent mean ± SEM. (d) Live-cell imaging of caspase-3/7 activity after 72-hour single-agent or combination treatments using a fluorescent apoptosis dye in isogenic IPM-BO-053 models. (e) Immunoblot analysis of cleaved PARP following 72-hour single-agent or combination treatments, with quantification of cleaved PARP/PARP ratios normalized to vehicle-treated controls. Error bars represent mean ± SEM. Representative results are shown from three independent experiments (n = 3). Statistical significance was determined using two-way ANOVA; *p < 0.05.

**Extended Data 6.**
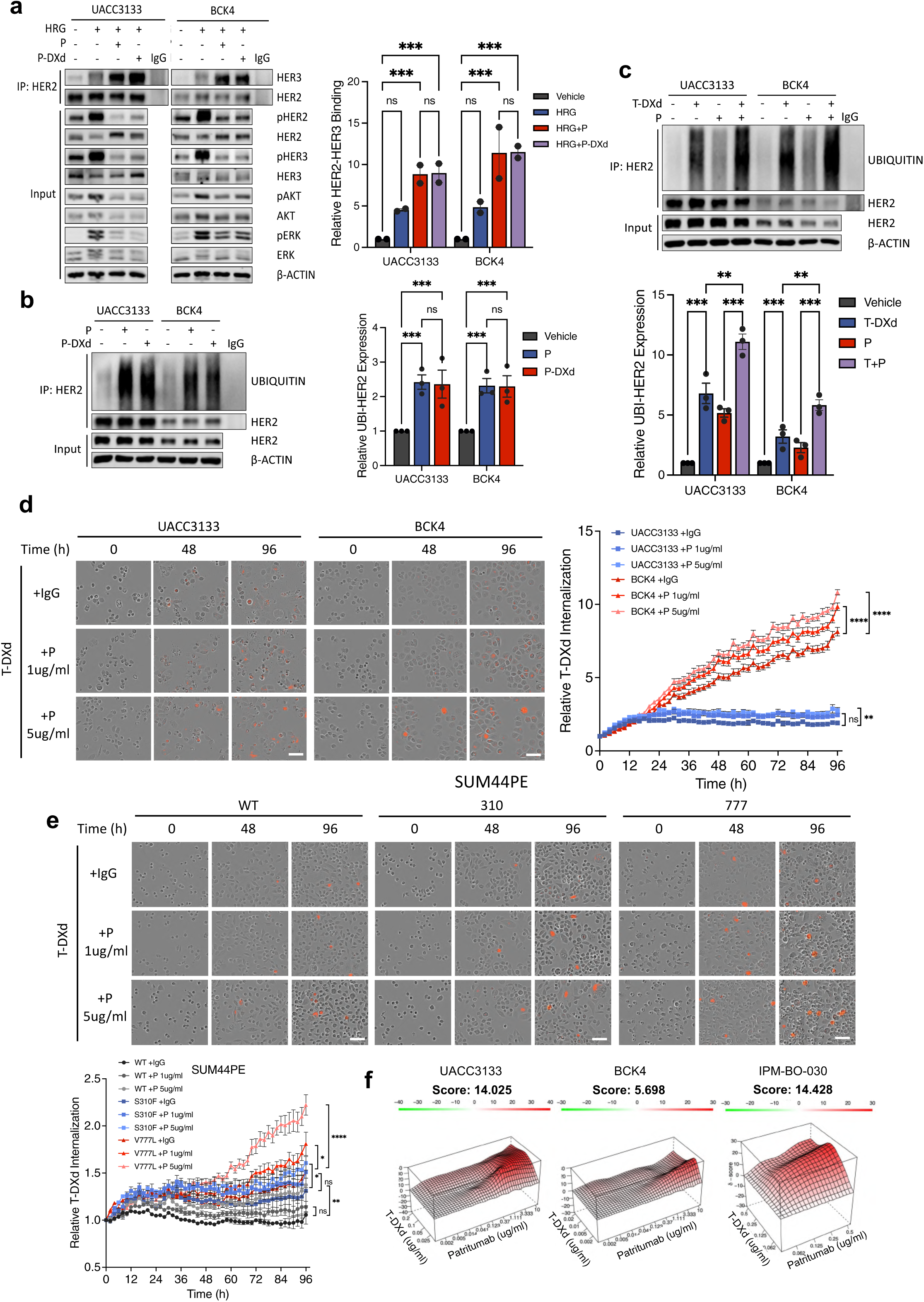
Patritumab induces HER2/HER3 interaction and HER2 ubiquitination and synergizes with T-DXd. (a) Immunoblot analysis of HER2/HER3 association and signaling inhibition in serum-starved UACC3133 and BCK4 treated with vehicle, HRG (1 nM), P-DXd (1 µg/mL), patritumab (1 µg/mL), or the combination for 10 minutes, with quantification normalized to vehicle-treated controls. Error bars represent mean ± SEM. (b) Immunoblot analysis of HER2 ubiquitination in serum-starved UACC3133 and BCK4 pretreated with bafilomycin A1 (100 nM, 2 h) followed by vehicle, P-DXd (1 µg/mL, 4 h), or patritumab (1 µg/mL, 4 h), with quantification normalized to vehicle-treated controls. Error bars represent mean ± SEM. (c) Immunoblot analysis of HER2 ubiquitination in serum-starved UACC3133 and BCK4 pretreated with bafilomycin A1 (100 nM, 2 h) followed by T-DXd (1 µg/mL, 4 h), patritumab (1 µg/mL, 4 h), or the combination. (d) Live-cell imaging of pHrodo-T-DXd internalization in UACC3133 and BCK4 treated with T-DXd (1 µg/mL) alone or with patritumab (1 or 5 µg/mL), with quantification over time normalized to baseline (t = 0) and confluency. Error bars represent mean ± SD. (e) Live-cell imaging of pHrodo-T-DXd internalization in isogenic SUM44PE models treated with T-DXd (1 µg/mL) alone or with patritumab (1 or 5 µg/mL), with quantification over time normalized to baseline (t = 0) and confluency. Error bars represent mean ± SD. (f) Dose-response analysis of combinatorial effects of T-DXd and patritumab in UACC3133, BCK4, and IPM-BO-030. Synergy scores were calculated using the HSA model; values > 10 indicate synergy. Representative results are shown from two to three independent experiments (n = 2-3). Statistical significance was determined using two-way ANOVA; *p < 0.05.

**Extended Data 7.**
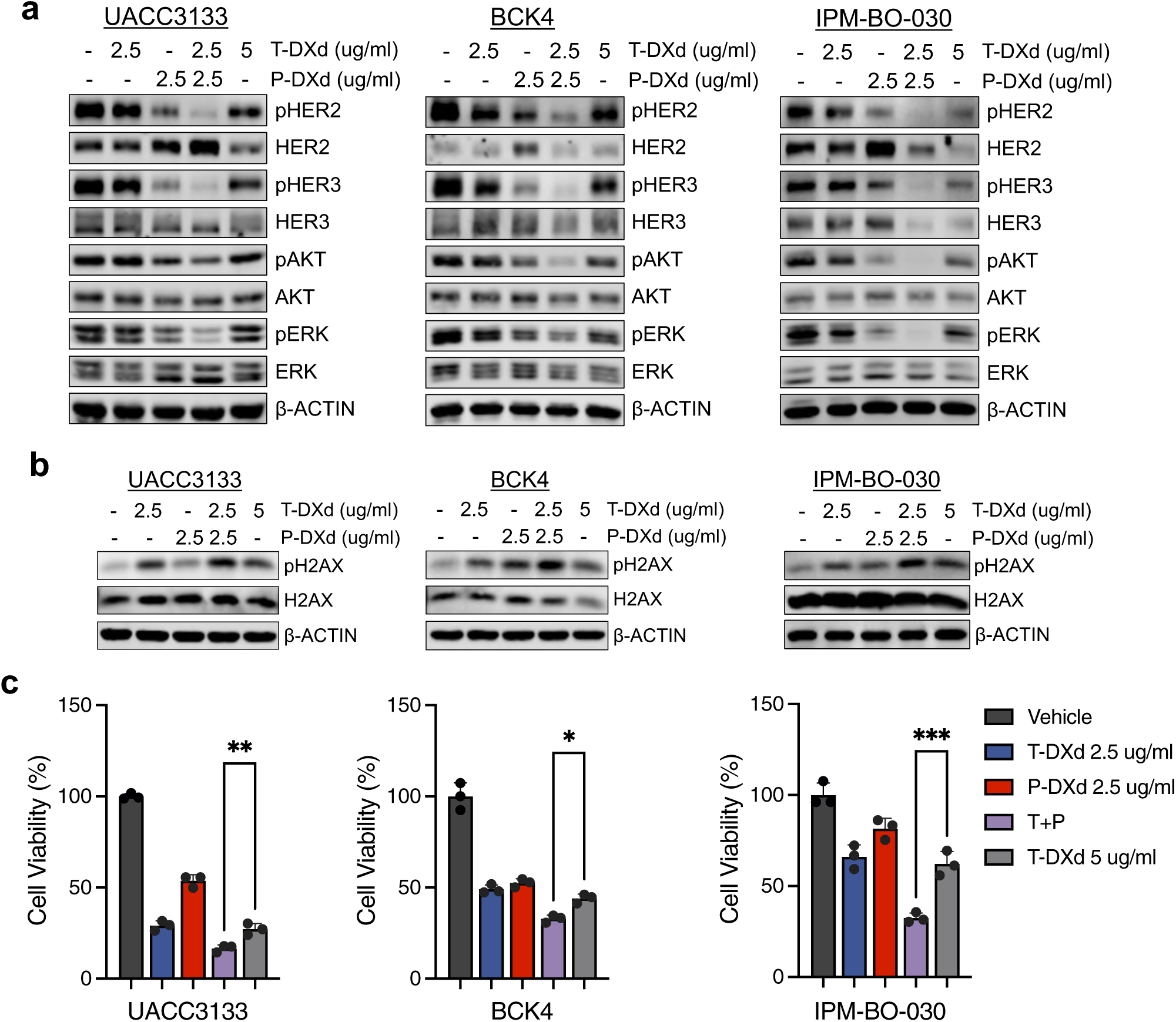
*In vitro* evaluation of T-DXd and P-DXd combination versus dose-equivalent T-DXd monotherapy in HER2-mutant ILC cell models. (a) Immunoblot analysis of HER2/HER3 and downstream signaling inhibition in serum-starved UACC3133, BCK4, and IPM-BO-030 treated with vehicle, T-DXd (2.5 µg/mL), P-DXd (2.5 µg/mL), the combination (2.5 + 2.5 µg/mL), or dose-equivalent high-concentration T-DXd monotherapy (5 µg/mL) for 4 hours followed by HRG stimulation (1 nM, 30 min). (b) Immunoblot analysis of pH2AX and H2AX following 72-hour treatment with the indicated conditions. (c) Cell viability analysis following 7-day treatment with the indicated conditions. Error bars represent mean ± SD. Representative results are shown from three independent experiments (n = 3). Statistical significance was determined using one-way ANOVA; *p < 0.05.

**Extended Data 8.**
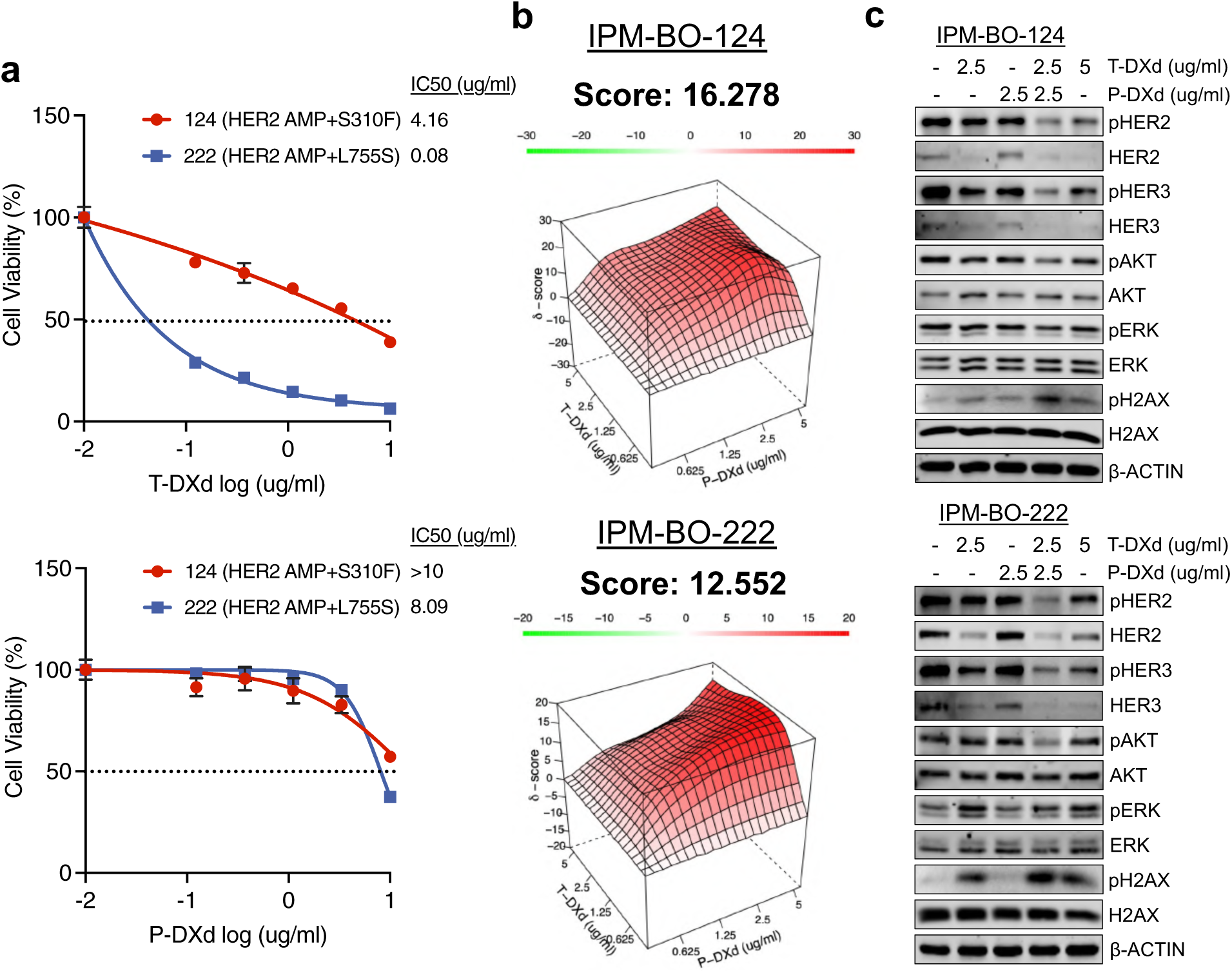
Evaluation of T-DXd and P-DXd combination therapy in PDOs with hotspot activating HER2 mutations. (a) Dose-response curves of IPM-BO-124 (HER2 AMP + S310F) and IPM-BO-222 (HER2 AMP + L755S) treated with T-DXd or P-DXd. (b) Dose-response analysis of the combinatorial effects of T-DXd and P-DXd in IPM-BO-124 and IPM-BO-222 PDOs. (c) Immunoblot analysis of HER2/HER3 and downstream signaling inhibition and DNA damage induction (pH2AX) following 72-hour treatment with vehicle, T-DXd (2.5 µg/mL), P-DXd (2.5 µg/mL), the combination (2.5 + 2.5 µg/mL), or dose-equivalent high-concentration T-DXd monotherapy (5 µg/mL). Representative results are shown from three independent experiments (n = 3). Synergy scores were calculated using the HSA model; values > 10 indicate synergy.

**Extended Data 9.**
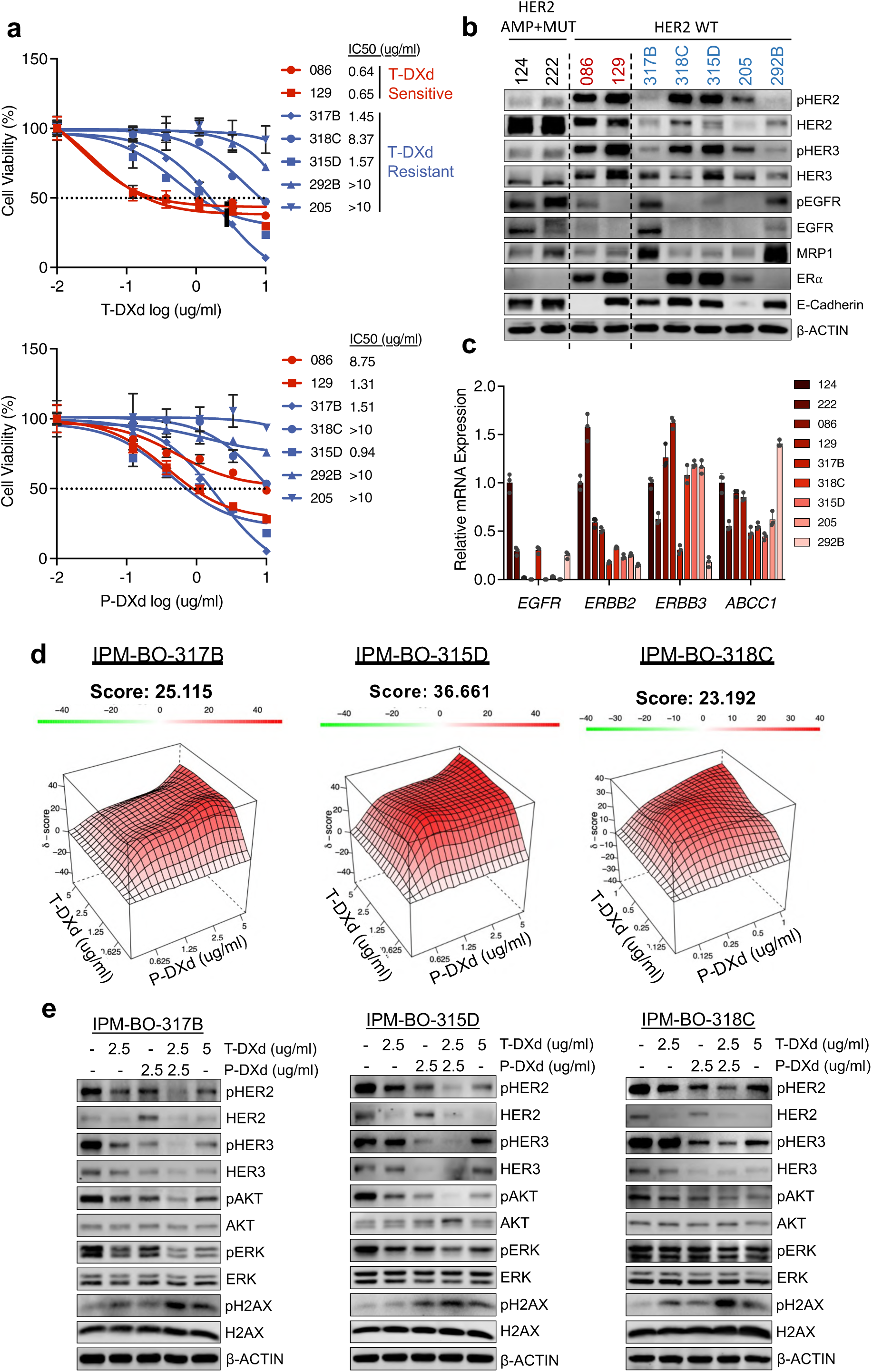
Evaluation of T-DXd and P-DXd response and molecular correlates of resistance across PDO models. (a) Dose-response curves of T-DXd or P-DXd across a panel of PDO models. Error bars represent mean ± SD. (b) Immunoblot analysis of EGFR, HER2, HER3, and MRP1 expression and activation across the PDO panel. (c) qRT-PCR analysis of EGFR (*EGFR*), HER2 (*ERBB2*), HER3 (*ERBB3*), and MRP1 (*ABCC1*) mRNA expression across the PDO panel. Error bars represent mean ± SD. Representative results are shown from three independent experiments (n = 3). (d) Dose-response analysis of combinatorial effects of T-DXd and P-DXd in T-DXd-resistant PDOs (IPM-BO-317B, IPM-BO-318C, IPM-BO-315D). Error bars represent mean ± SD. (e) Immunoblot analysis of HER2/HER3 and downstream signaling inhibition and DNA damage induction (pH2AX) following 72-hour treatment with vehicle, T-DXd (2.5 µg/mL), P-DXd (2.5 µg/mL), the combination (2.5 + 2.5 µg/mL), or dose-equivalent high-concentration T-DXd monotherapy (5 µg/mL). Representative results are shown from three independent experiments (n = 3). Synergy scores were calculated using the HSA model; values > 10 indicate synergy.

**Extended Data 10.**
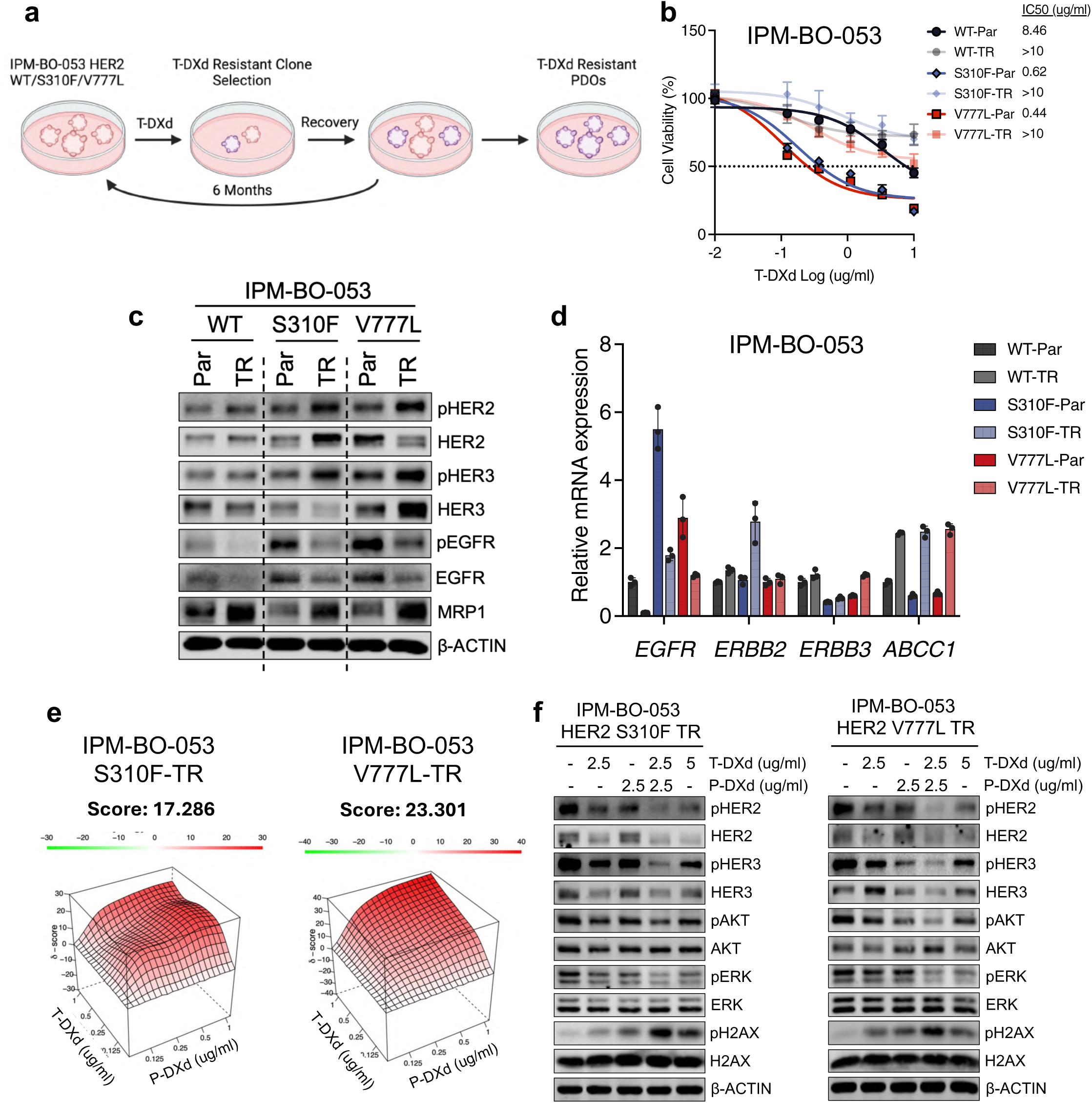
Characterization of acquired T-DXd resistance in isogenic HER2-wild-type and HER2-mutant PDOs. (a) Schematic depicting generation of acquired T-DXd-resistant IPM-BO-053 PDOs. (b) Dose-response curves of T-DXd treatment in parental (Par) and acquired T-DXd-resistant (TR) isogenic IPM-BO-053 models. Error bars represent mean ± SD. (c) Immunoblot analysis of EGFR, HER2, HER3, and MRP1 protein expression and activation in parental and resistant models. (d) qRT-PCR analysis of EGFR (*EGFR*), HER2 (*ERBB2*), HER3 (*ERBB3*), and MRP1 (*ABCC1*) mRNA expression in parental and resistant models. Error bars represent mean ± SD. (e) Dose-response analysis of combinatorial effects of T-DXd and P-DXd in acquired T-DXd-resistant IPM-BO-053 S310F and IPM-BO-053 V777L models. Error bars represent mean ± SD. (f) Immunoblot analysis of HER2/HER3 and downstream signaling inhibition and DNA damage induction (pH2AX) following 72-hour treatment with vehicle, T-DXd (2.5 µg/mL), P-DXd (2.5 µg/mL), the combination (2.5 + 2.5 µg/mL), or dose-equivalent high-concentration T-DXd monotherapy (5 µg/mL). Representative results are shown from three independent experiments (n = 3). Synergy scores were calculated using the HSA model; values > 10 indicate synergy.

**Supplementary Figure 1.**
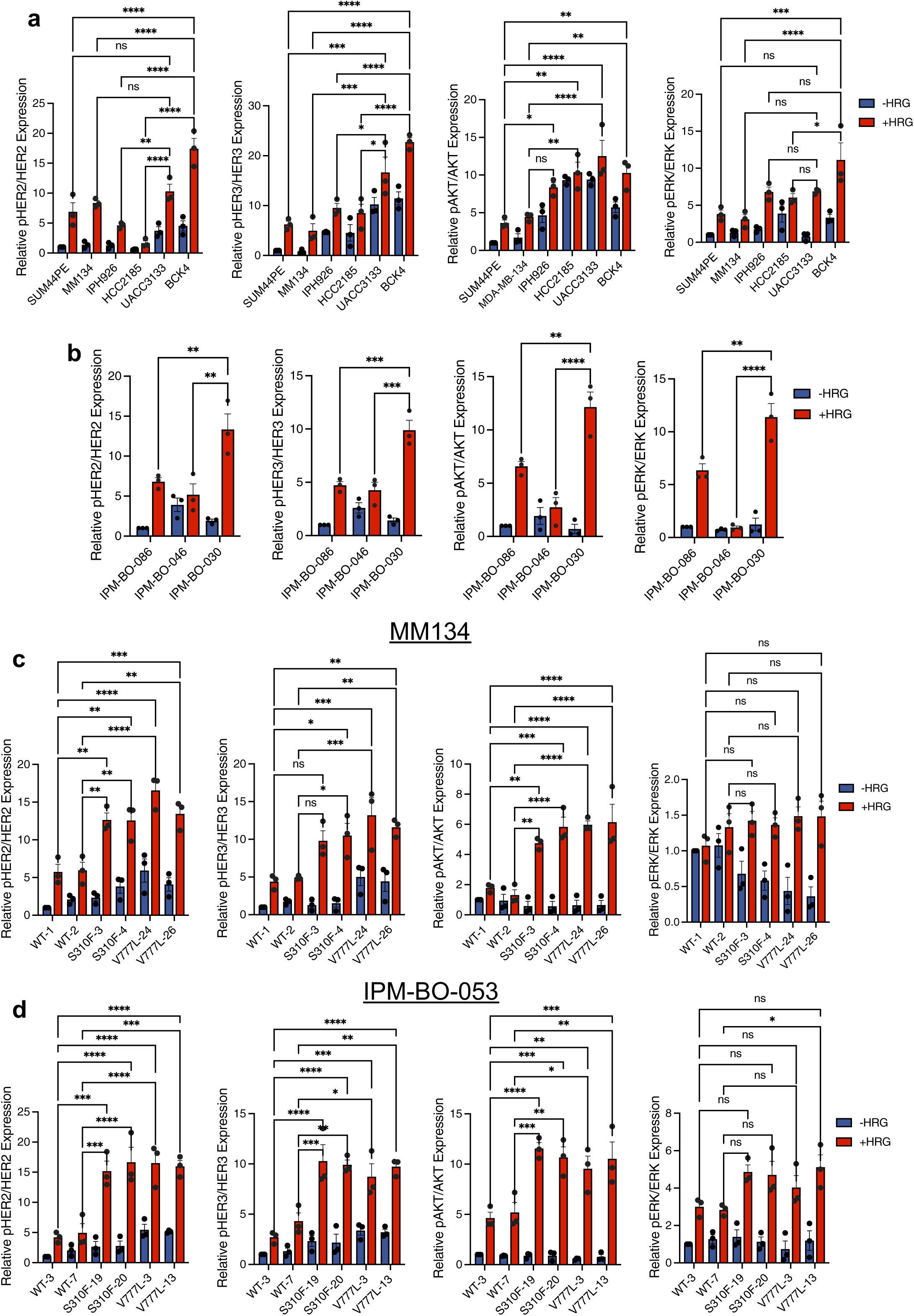
Quantification of signaling activation in HER2-mutant and wild-type ILC models. (a-d) Quantification of phospho-to-total protein ratios for HER2, HER3, AKT, and ERK normalized to wild-type controls in endogenous cell line models (a), endogenous PDO models (b), isogenic MM134 models (c), and isogenic IPM-BO-053 models (d). Error bars represent mean ± SEM. Representative results are shown from three independent experiments (n = 3). Statistical significance was determined using two-way ANOVA; *p* < 0.05.

**Supplementary Figure 2.**
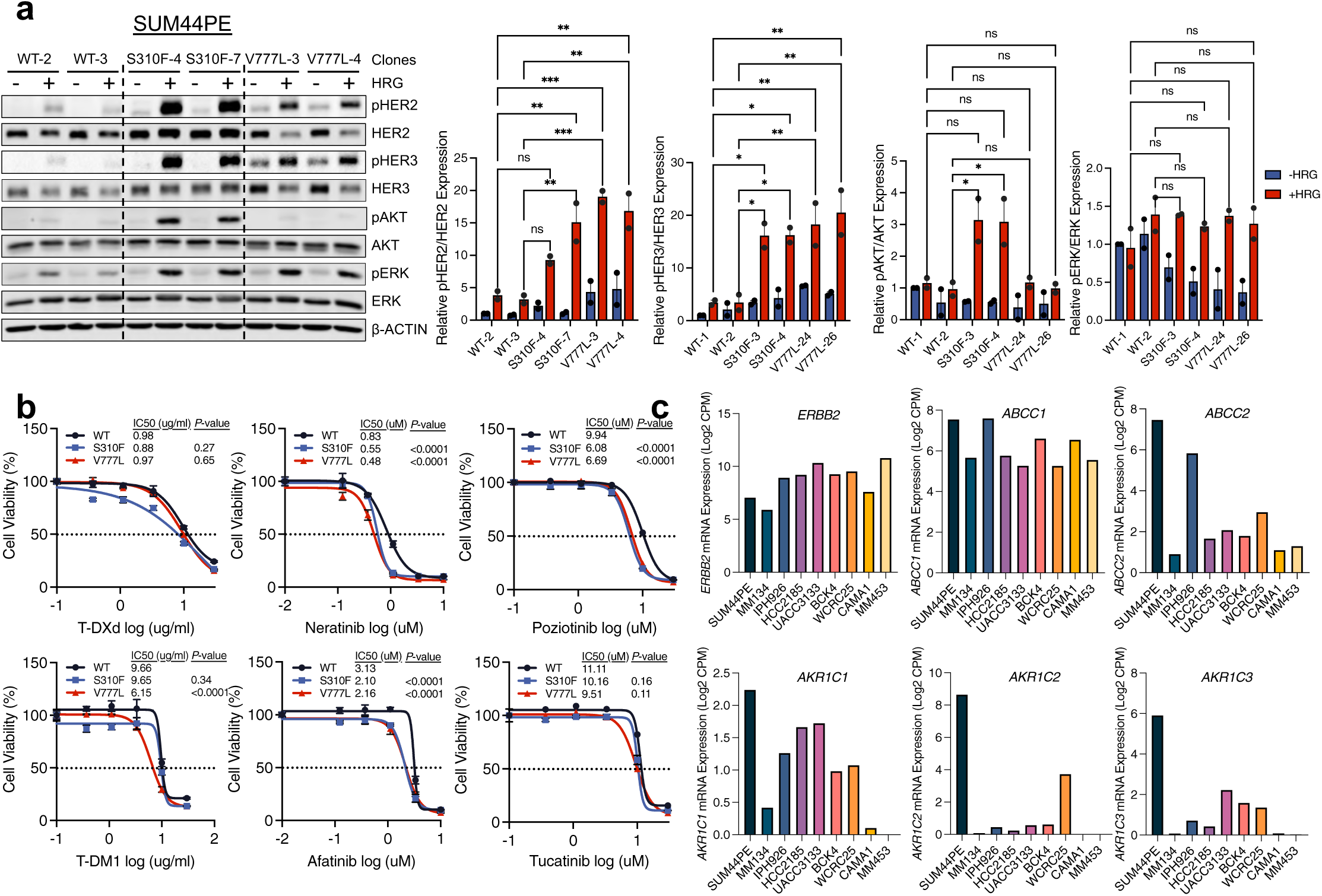
Characterization of SUM44PE HER2 wild-type and mutant cells. (a) Immunoblot analysis of HER2 and downstream signaling under serum-starved and HRG-stimulated conditions (1 nM, 10 min) in SUM44PE isogenic HER2 wild-type and mutant clones, with quantification of phospho-to-total protein ratios (HER2, HER3, AKT, ERK) normalized to wild-type controls. Error bars represent mean ± SEM. Statistical significance was determined using two-way ANOVA; *p < 0.05. (b) Dose-response curves comparing sensitivity of isogenic SUM44PE models to HER2-targeted TKIs and ADCs. Error bars represent mean ± SD. Statistical significance was determined using sum-of-squares global f-test; *p < 0.05. (c) Transcriptomic profiling of genes associated with established ADC-resistance mechanisms across ILC cell lines. Representative results are shown from two to three independent experiments (n = 2-3).

**Supplementary Figure 3.**
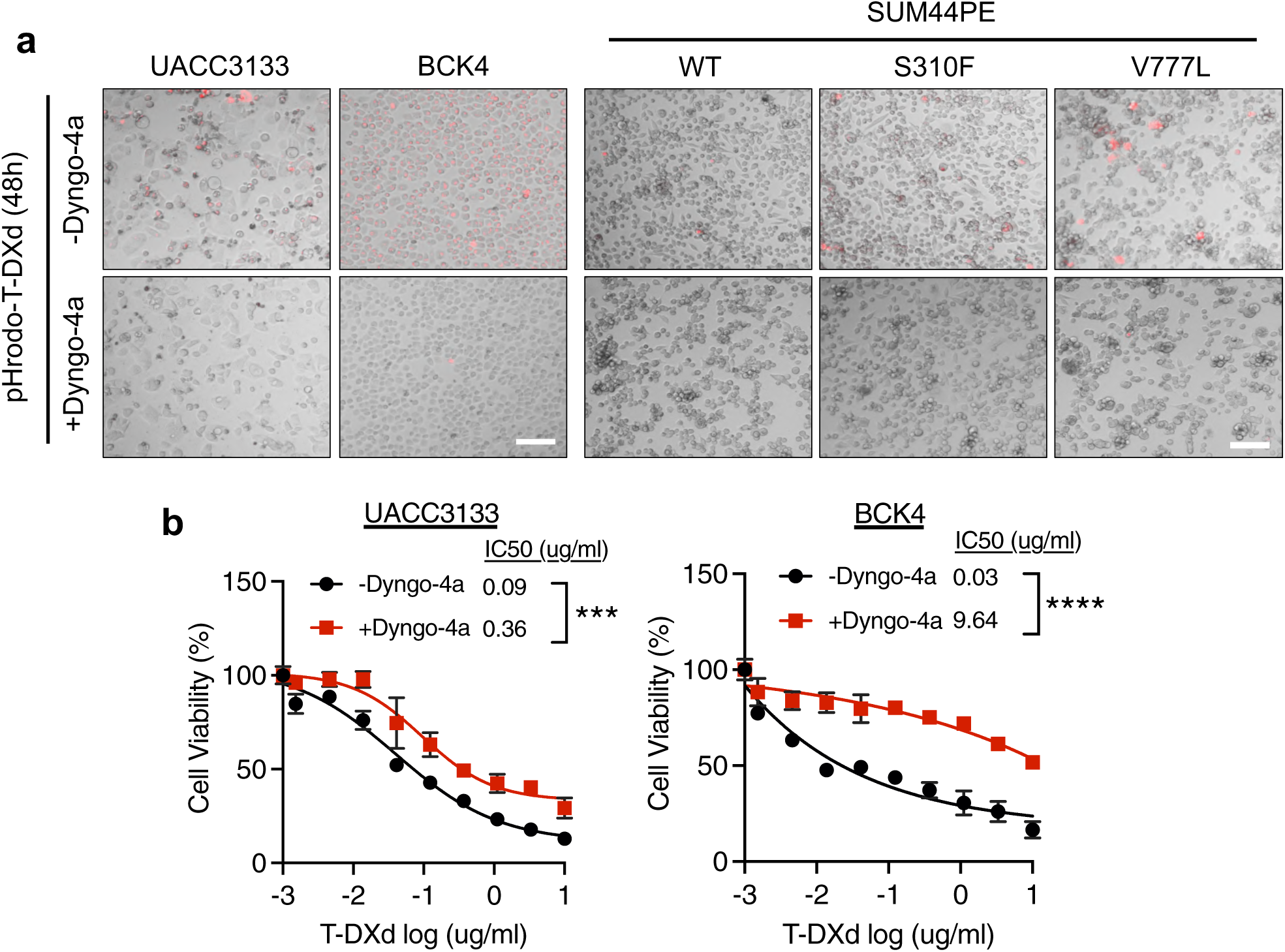
Dynamin inhibition blunts T-DXd internalization and cytotoxicity. (a) Live-cell imaging of pHrodo-T-DXd internalization in UACC3133 and BCK4 and isogenic SUM44PE models pretreated with vehicle or the dynamin inhibitor Dyngo-4a (10 µM, 1 h). (b) Dose-response curves demonstrating attenuation of T-DXd cytotoxicity by Dyngo-4a co-treatment in UACC3133 and BCK4. Error bars represent mean ± SD. Representative results are shown from three independent experiments (n = 3). Statistical significance was determined using a sum-of-squares Global F-test; *p < 0.05. Scale bar, 100 µm.

**Supplementary Figure 4.**
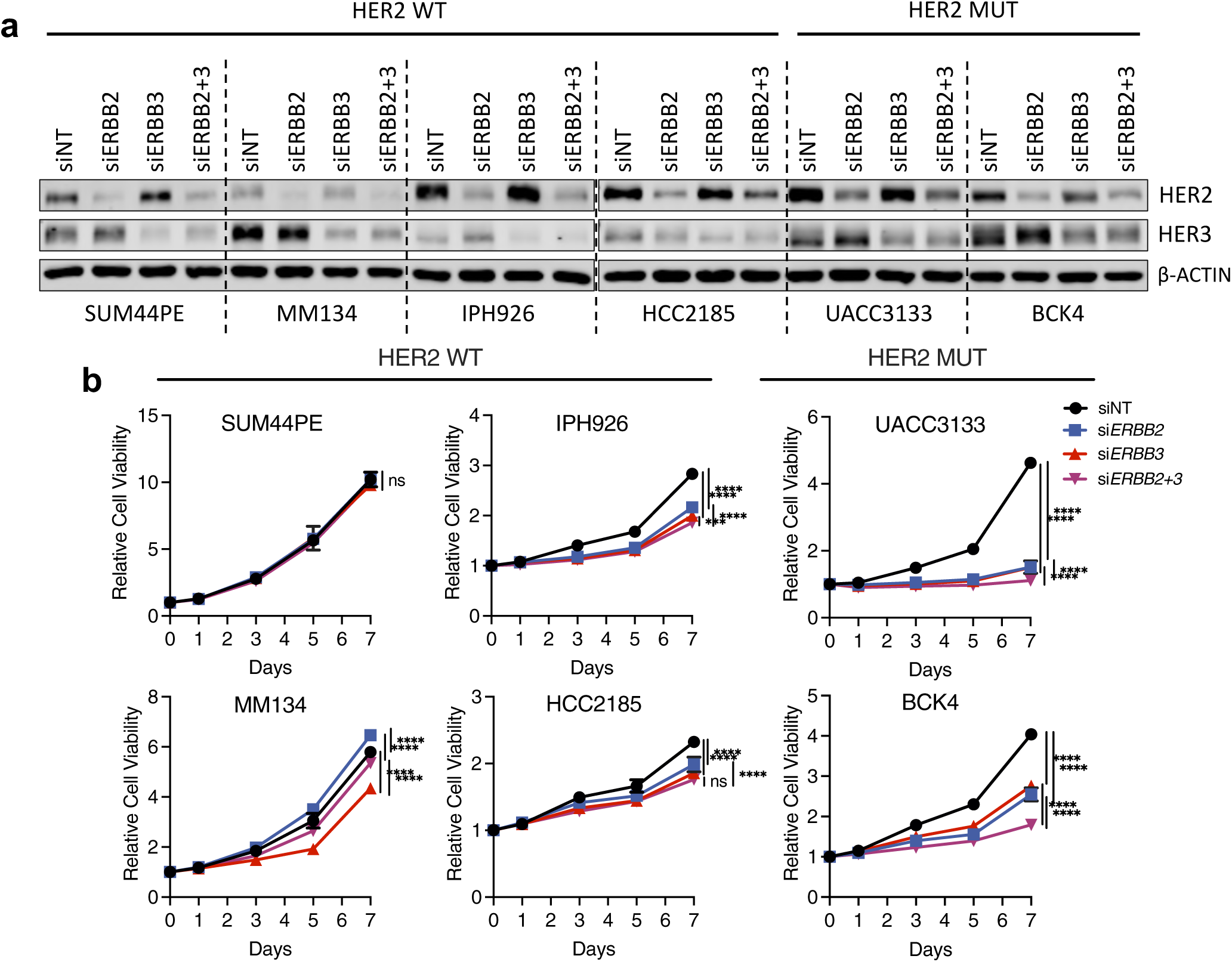
HER2-mutant ILC cells exhibit dual dependence on HER2 and HER3 for growth. (a) Immunoblot analysis of siRNA-mediated knockdown of HER2, HER3, or both in endogenous HER2-wild-type (SUM44PE, MM134, IPH926, HCC2185) and HER2-mutant (UACC3133, BCK4) ILC cell lines. (b) Quantification of cell viability following single or dual receptor silencing across the indicated ILC cell lines. Representative results are shown from three independent experiments (n = 3). Error bars represent mean ± SD. Statistical significance was determined using two-way ANOVA; *p < 0.05.

**Supplementary Figure 5.**
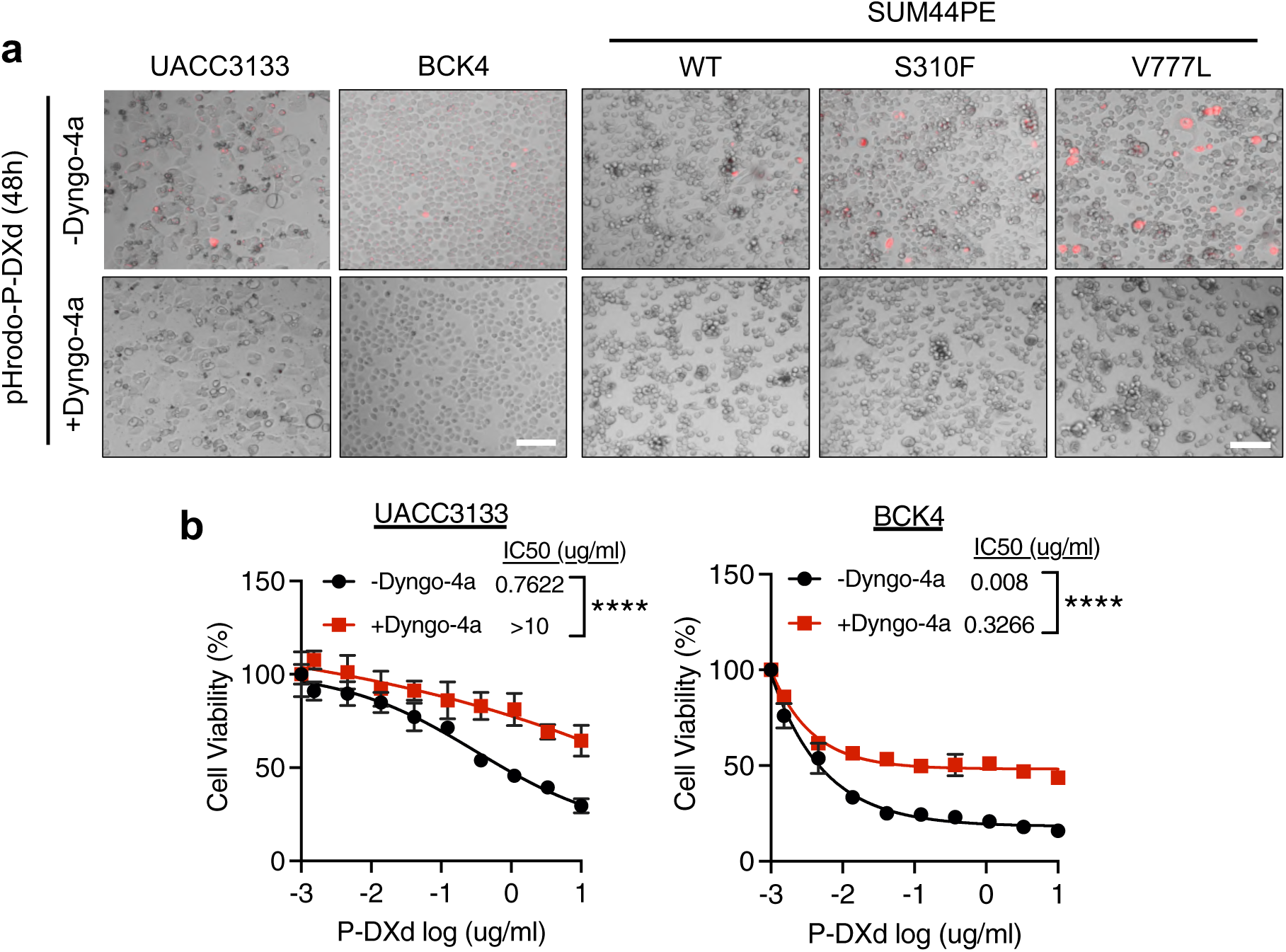
Dynamin inhibition blunts P-DXd internalization and cytotoxicity. (a) Live-cell imaging of pHrodo-P-DXd internalization in UACC3133, BCK4 and isogenic SUM44PE models pretreated with vehicle or Dyngo-4a (10 µM, 1 h).b) Dose-response curves demonstrating attenuation of P-DXd cytotoxicity by Dyngo-4a co-treatment in UACC3133 and BCK4. Error bars represent mean ± SD. Representative results are shown from three independent experiments (n = 3). Statistical significance was determined using a sum-of-squares Global F-test; *p < 0.05. Scale bar, 100 µm.

**Supplementary Figure 6.**
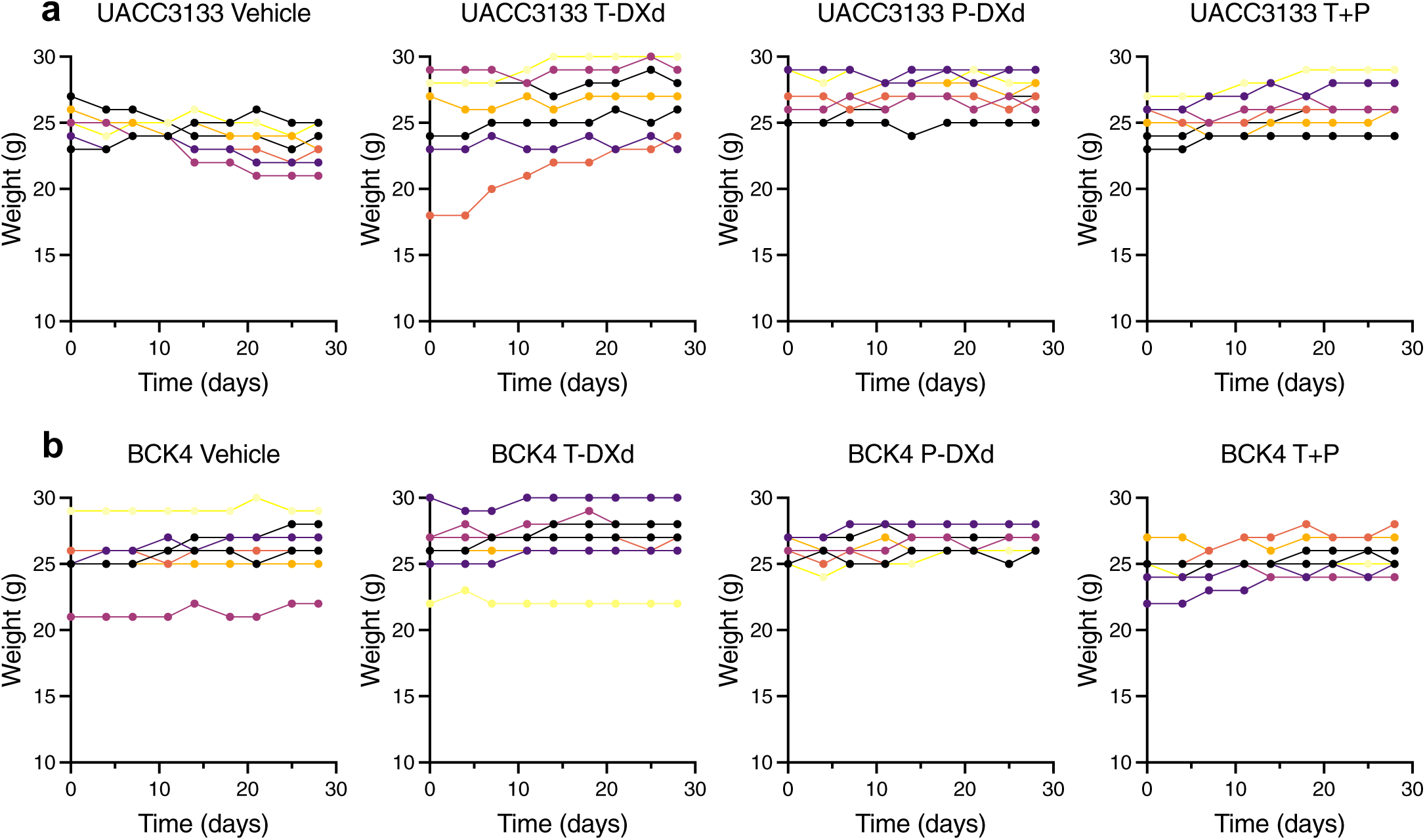
Body weight assessment of ADC-associated toxicity during T-DXd, P-DXd, and combination treatment in HER2-mutant ILC xenografts. (a) Body weight monitoring of UACC3133 xenografts treated with vehicle, T-DXd (2.5 mg/kg), P-DXd (2.5 mg/kg), or the combination (T + P; 5 mg/kg total) over a 28-day treatment period. (b) Body weight monitoring of BCK4 xenografts treated with vehicle, T-DXd (2.5 mg/kg), P-DXd (2.5 mg/kg), or the combination (T + P; 5 mg/kg total) over a 28-day treatment period.

**Supplementary Figure 7.**
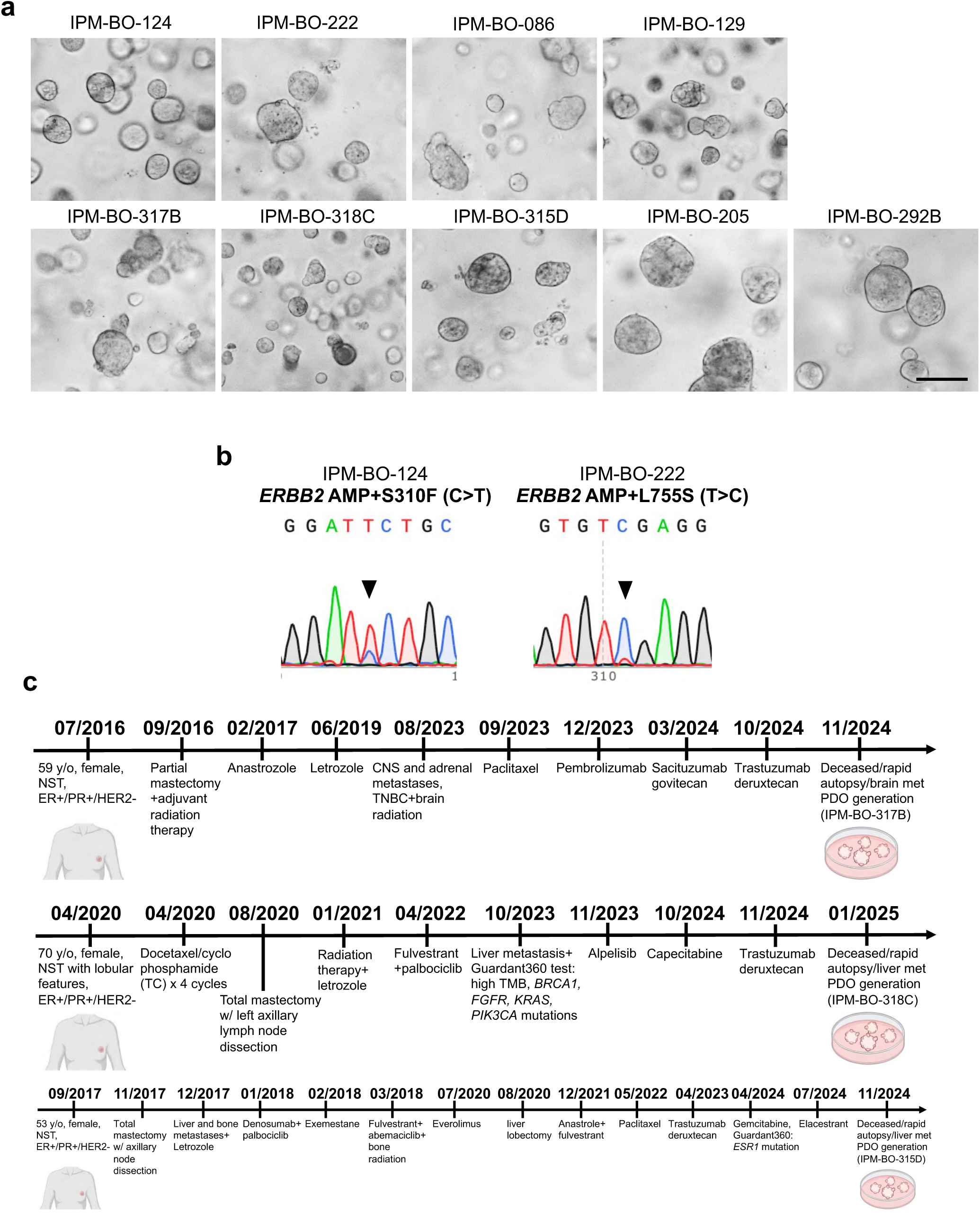
PDO characteristics. (a) Bright-field images illustrating the morphological diversity of PDOs used in this study. Scale bar, 300 µm. (b) Sanger sequencing chromatograms confirming hotspot HER2 mutations S310F and L755S in PDO models IPM-BO-124 and IPM-BO-222, respectively. (c) Schematic representation of treatment histories for patients from whom T-DXd-resistant PDOs (IPM-BO-317B, IPM-BO-318C, IPM-BO-315D) were generated. Intrinsic resistance: IPM-BO-317B, IPM-BO-318C; acquired resistance: IPM-BO-315D.

**Supplementary Table 1.** PDO characteristics. Characteristics of patient-derived organoids (PDOs) used in this study.

| PDO ID | Tumor Histology Subtype | Tumor Origin | Metastatic Site | Tumor ER/HER2 Status | T-DXd Therapy (Y/N) |
| --- | --- | --- | --- | --- | --- |
| IPM-BO-124 | NST | Primary | N/A | ER-/HER2 AMP+S310F | N |
| IPM-BO-222 | NST | Primary | N/A | ER-/HER2 AMP+L755S | N |
| IPM-BO-086 | ILC | Metastatic | Bone | ER+/HER2- | N |
| IPM-BO-129 | NST | Metastatic | Bone | ER+/HER2- | N |
| IPM-BO-317B | NST | Metastatic | Brain | ER-/HER2- | Y (progression after 1 month) |
| IPM-BO-318C | NST w/ lobular features | Metastatic | Liver | ER+/HER2- | Y (progression after 2 months) |
| IPM-BO-315D | NST | Metastatic | Liver | ER+/HER2- | Y (progression after 1 year) |
| IPM-BO-205 | NST | Metastatic | Ascites | ER+/HER2- | N |
| IPM-BO-292B | NST | Metastatic | Lung | ER+/HER2- | N |

## Notes

### Summary of Updates

I accidentally placed my PI (Steffi Oesterreich) as the first author when filling out the author list. This error has been corrected. The actual manuscript pdf has the correct order.

